# Alternative splicing expands and remodels the breast cancer proteoform landscape

**DOI:** 10.64898/2026.04.30.722115

**Authors:** Felicia T. Jiang, Dengwang Chen, Ziwei Wang, Xiangeng Wang, Hao Hao, Yinghan Zhang, Runhao Zhao, Feng Liang, Taoyong Cui, Zhenchao Tang, Tianli Luo, Yi Shuai, Minghao Xu, Chenchen Xu, Jia Xu, Wentao Zhang, Jun Tan, Xin Wang

## Abstract

Alternative splicing generates extensive transcript diversity, but the extent to which this diversity contributes to the translated proteome and reshapes protein function remains unclear. Here, we develop 3DisoGalaxy, an atlas integrating transcript reconstruction, ribosome profiling, structural modelling and regulatory annotation in breast cancer. We find that 58,292 of 90,929 transcripts are linked to translation-supported ORFs, 71.9% of which arise from non-canonical transcript structures. These products are broadly accommodated within established protein fold space, indicating that alternative isoforms frequently preserve recognizable protein architectures despite substantial sequence variation. Functional divergence is instead concentrated in domains, intrinsically disordered regions, localization determinants and regulatory motifs. Motif-repertoire remodelling is more extensive in cancer-driver genes than in other genes, linking isoform-associated regulatory variation to cancer-relevant proteins. As an example of domain remodelling, we identify an AKT1-ΔPH isoform that lacks the N-terminal PH domain while retaining the kinase core, with immunoblotting revealing a lower-molecular-weight AKT1 species consistent with this isoform. Together, our results show that alternative transcript diversity extensively enters translation-supported proteoform space and establish a systematic link between transcript variation and protein functional diversification.

**Graphical Abstract:** 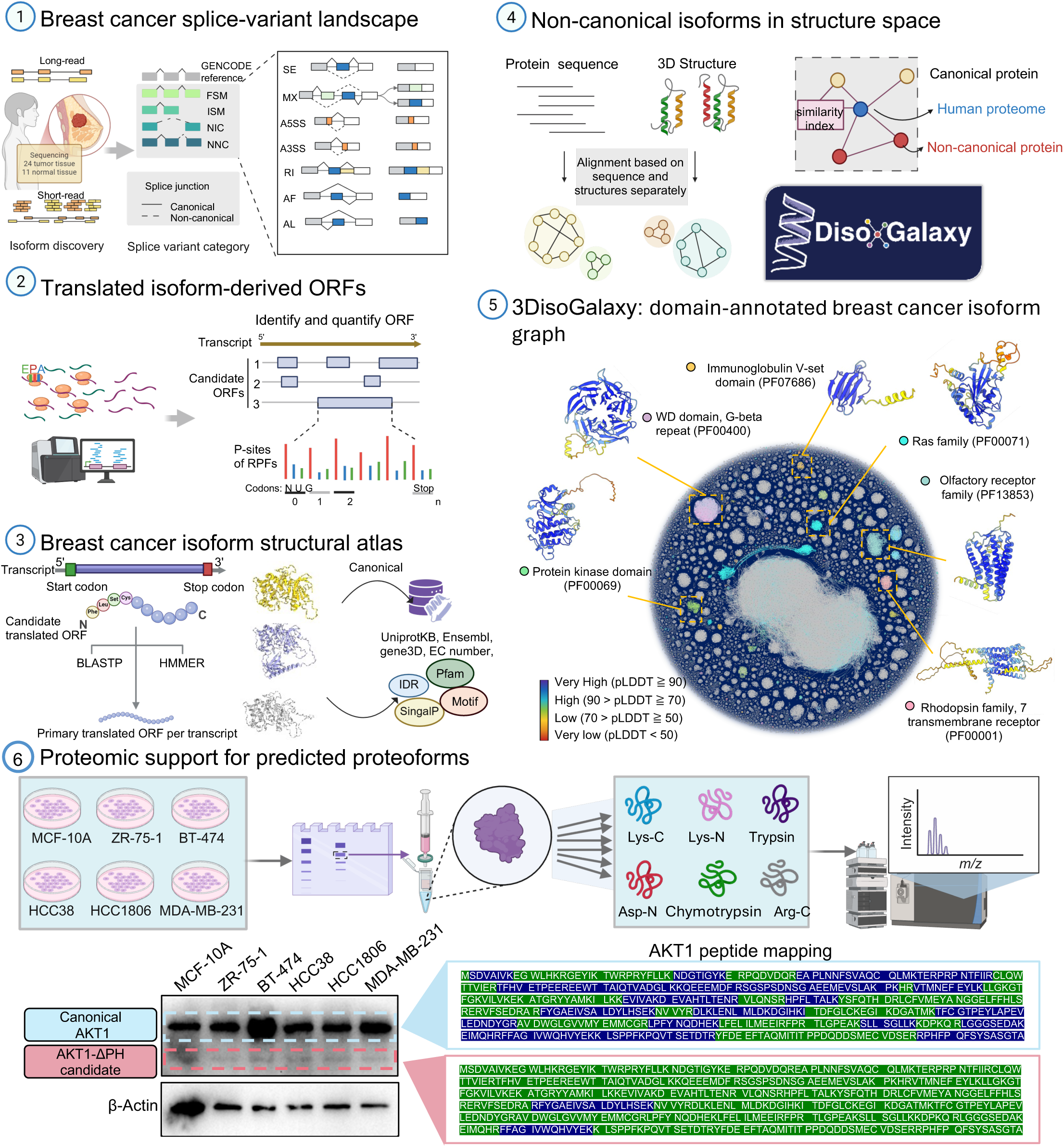

## Introduction

Alternative RNA processing is a major source of molecular diversity in human cells, allowing individual genes to generate multiple full-length transcripts with distinct coding sequences, untranslated regions and regulatory contexts. Genome-wide studies established that alternative splicing affects most human multiexon genes and is extensively regulated across tissues and cell types (Wang et al., 2008; Pan et al., 2008). The resulting protein products form one component of the broader proteoform diversity arising from genetic variation, RNA processing and post-translational modification (Smith and Kelleher, 2013; Aebersold et al., 2018). Alternative protein isoforms need not behave as minor variants of a single canonical product. Tissue-specific splicing can alter disordered segments containing interaction motifs, and systematic interactome profiling has shown that isoforms encoded by the same gene frequently engage substantially different interaction partners (Buljan et al., 2012; Yang et al., 2016). This variation is particularly relevant in cancer, where recurrent alterations in splicing factors and widespread tumour-associated splicing changes contribute an additional layer of molecular heterogeneity beyond gene expression and somatic mutation (Dvinge et al., 2016; Kahles et al., 2018). Alternative transcripts may therefore encode substantial protein-level diversity that cannot be resolved through gene-level analysis alone.

The extent to which this transcript diversity contributes to the proteome remains uncertain. Long-read RNA sequencing can resolve complete transcript architectures that are difficult to reconstruct from fragmented short-read data and has revealed extensive previously unannotated transcript diversity across human cells and tissues (Sharon et al., 2013; Glinos et al., 2022). Detection of a full-length transcript, however, does not establish that it is productively translated or gives rise to a detectable protein. Conventional bottom-up mass spectrometry also has limited power to distinguish closely related proteoforms because most proteolytic peptides are shared among isoforms, whereas the few isoform-specific regions may not produce detectable peptides or may occur in proteins with low or context-dependent abundance (Aebersold and Mann, 2016). Integrated transcriptomic and proteomic analyses have shown that changes in transcript usage can propagate to the protein level, and tissue-informed proteogenomic searches have identified non-canonical protein isoforms that are absent from standard protein databases (Liu et al., 2017; Lau et al., 2019). Nevertheless, the number of isoforms resolved by unique peptide evidence remains small relative to the diversity observed by transcript sequencing. This discrepancy may reflect both biological non-productivity of some alternative transcripts and systematic under-detection of isoform-specific protein products by current proteomic technologies. Determining how much full-length alternative transcript diversity enters the translated proteoform repertoire therefore remains a central problem.

Ribosome profiling provides an independent route for addressing this problem by sequencing ribosome-protected RNA fragments and resolving translated open reading frames at near-nucleotide resolution (Ingolia et al., 2009). Isoform-level analyses have shown that a substantial fraction of expressed transcript isoforms is associated with translation and that changes in transcript usage can produce concordant changes in ribosome occupancy (Reixachs-Solé et al., 2020). Translation support alone, however, does not establish that an open reading frame encodes a structurally coherent and biologically interpretable protein form. A ribosome-associated sequence may lack a recognizable folded core or fail to exhibit the expected relationships among sequence composition, structural organization, cellular localization and functional annotation. Assessing proteoform coherence therefore requires convergent evidence that translated sequences follow established protein-level principles. Advances in protein structure prediction now permit structural analysis of sequences without experimentally determined models, including predictions at the scale of nearly the entire human proteome (Jumper et al., 2021; Tunyasuvunakool et al., 2021). These structural models can be considered alongside intrinsic disorder, subcellular localization and functional annotation, properties that are independently linked to protein organization within the cell (Thul et al., 2017). No individual prediction establishes protein function, but concordance across these properties can reveal whether a translated sequence repertoire behaves as an organized protein space rather than an arbitrary collection of open reading frames.

Biologically coherent proteoforms may acquire distinct regulatory properties without generating entirely new protein folds. Alternative exon usage can modify terminal regions, domain composition, intrinsically disordered segments and localization-associated sequences while often preserving major folded regions. Intrinsically disordered regions are particularly enriched for short interaction and regulatory motifs, and tissue-specific splicing of these regions can rewire protein interaction networks (Nguyen Ba et al., 2012; Buljan et al., 2012). Proteogenomic analyses have similarly shown that alternatively spliced protein regions frequently overlap disordered sequences and post-translational modification sites (Lau et al., 2019). The gain, loss or repositioning of short regulatory motifs could therefore alter interaction specificity, cellular targeting, protein stability or signalling without globally disrupting protein architecture. This model is consistent with experimental interaction maps in which isoforms from the same gene preserve portions of a common interaction repertoire while acquiring or losing isoform-specific partners (Yang et al., 2016). Transcript reconstruction, translation analysis, structure prediction and regulatory annotation have nevertheless largely been investigated separately. An integrated disease-scale framework is needed to determine how extensively transcript diversity enters protein space, whether the resulting proteoforms exhibit coherent structural and cellular organization, and how related proteoforms acquire regulatory diversity.

Here, we establish an integrative computational framework to determine how full-length transcript diversity propagates into translation-supported protein space and how the resulting protein isoforms diversify structurally and functionally. We implement this framework through 3DisoGalaxy, a structure-function inference atlas of breast cancer proteoforms. Long-read transcript reconstruction and independent short-read evidence were combined to define complete, evidence-supported transcript sequences. Ribosome profiling was then used to assign translation-supported open reading frames, followed by AlphaFold-based structure prediction and analyses of intrinsic disorder, subcellular localization, protein domains and regulatory motifs. The resulting structures were organized within a structure-similarity network, allowing isoform-derived proteins to be interpreted relative to canonical isoforms, representative human proteome structures and established protein families. This design addressed three linked questions: how much full-length alternative transcript diversity enters the translation-supported proteoform repertoire, whether the resulting proteoforms exhibit coherent protein-level organization, and which molecular features are remodeled as related proteoforms diversify.

## Results

### Hybrid transcriptomics resolves an extensive and evidence-supported breast cancer transcript repertoire

To establish a sequence-resolved foundation for proteoform analysis, we integrated PacBio long-read RNA sequencing from 35 breast tissue samples, short-read RNA-seq from four independent breast cancer cohorts comprising 339 samples, and ribosome profiling from 42 breast epithelial and breast cancer cell-line samples. These datasets formed the basis of 3DisoGalaxy, an interactive framework linking full-length transcript reconstruction with translation support, predicted protein structure and regulatory annotation **(Supplementary Fig. 1; Supplementary Fig. 2a; Supplementary Table 1)**.

Full-length transcript models were reconstructed from 24 breast tumours and 11 adjacent normal tissues and classified by SQANTI3 as full-splice matches (FSMs), incomplete-splice matches (ISMs), novel-in-catalogue (NIC) transcripts, novel-not-in-catalogue (NNC) transcripts or other structural categories according to their splice-junction and transcript-boundary relationships with reference annotations **(Fig. 1a**; Pardo-Palacios et al., 2024). Their exon architectures were further resolved into major alternative splicing event classes, including skipped exons (SE), mutually exclusive exons (MXE), alternative 5′ and 3′ splice sites (A5SS and A3SS), retained introns (RI), and alternative first and last exons (AF and AL) (Alamancos et al., 2015). Independent RNA-seq, poly(A)-site and CAGE data were subsequently incorporated to support splice junctions and transcript boundaries before proteoform-level analysis.

**Figure 1.**
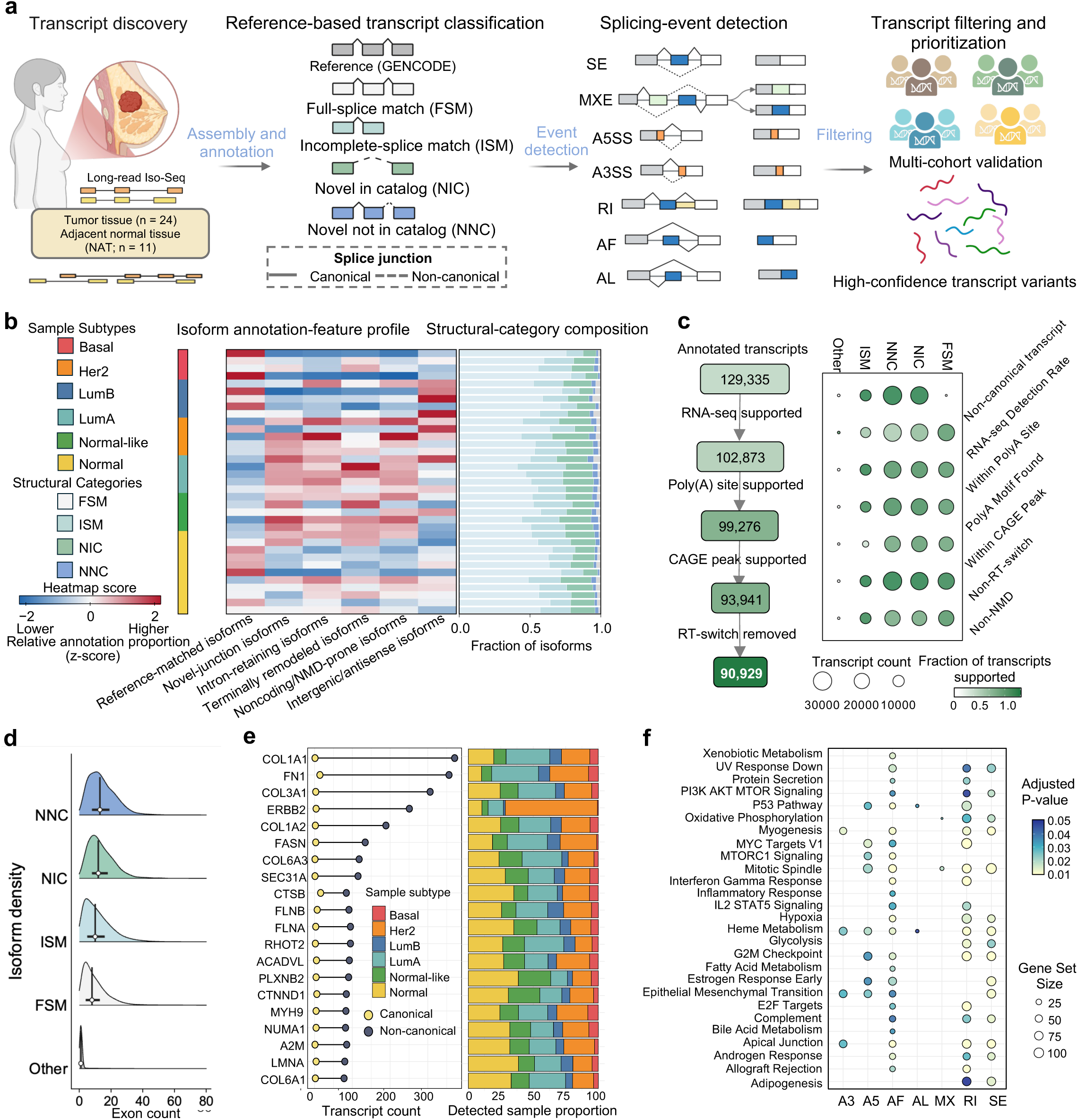
Hybrid transcriptomics defines a high-confidence splicing-variant landscape for breast cancer. **a**, Workflow integrating long-read Iso-Seq data from breast tumours (n = 24) and adjacent normal tissues (NAT; n = 11), reference-based transcript classification, alternative splicing-event detection, and multi-cohort transcript filtering and prioritization. **b,** Sample-level isoform annotation-feature profiles and transcript structural-category composition across breast cancer subtypes and normal tissues. Heatmap values represent relative annotation proportions expressed as z scores. **c,** Evidence-based transcript curation integrating short-read RNA-seq, poly(A)-site and CAGE-peak support, together with removal of reverse-transcription switching artefacts, reducing the initial set of 129,335 annotated transcripts to 90,929 high-confidence transcript variants. The accompanying matrix summarizes transcript counts and the fraction of transcripts meeting each annotation or quality-control criterion. **d,** Exon-count distributions across transcript structural categories. **e,** Top 20 genes ranked by transcript count, showing reference-matched and non-reference isoform contributions and subtype-resolved sample detection proportions. **f,** Pathway gene set enrichment of subtype-associated alternative splicing variants stratified by splicing-event type. Dot size denotes gene set size, and colour indicates the Benjamini–Hochberg-adjusted *P* value. FSM, full-splice match; ISM, incomplete-splice match; NIC, novel in catalog; NNC, novel not in catalog; SE, skipped exon; MXE, mutually exclusive exons; A5SS and A3SS, alternative 5′ and 3′ splice sites, respectively; RI, retained intron; AF, alternative first exon; AL, alternative last exon; CAGE, cap analysis of gene expression.

The long-read cohort comprised Basal (n = 4), HER2 (n = 6), Luminal A (n = 5), Luminal B (n = 4), Normal-like (n = 5) and adjacent normal tissue samples (n = 11) **(Supplementary Fig. 2b)**. Across the pooled transcript catalogue, NNC and NIC models accounted for 33.5% and 29.4% of transcript variants, respectively, compared with 26.6% FSM and 10.5% ISM models **(Supplementary Fig. 2c)**. Structurally novel NIC and NNC transcripts therefore comprised 62.9% of the reconstructed repertoire. The reconstructed transcript isoforms showed a predominantly unimodal length distribution, with most transcripts ranging from approximately 10³ to 10⁴ nucleotides (**Supplementary Fig. 2d**). At the individual-sample level, however, FSM transcripts generally remained the largest structural category, whereas ISM, NIC and NNC transcripts were detected across all sample groups **(Fig. 1b)**. Thus, structurally novel transcripts substantially expanded the pooled catalogue without dominating any single sample group.

Annotation-feature profiles also varied among individual samples. Several HER2, Luminal A and Normal-like samples showed relatively higher representation of novel-junction, intron-retaining, terminally remodelled and noncoding or nonsense-mediated-decay-prone isoforms, whereas reference-matched isoforms were more prominent in several Basal and Luminal B samples **(Fig. 1b)**. These patterns indicated sample-level heterogeneity in transcript architecture, although non-reference isoforms were broadly distributed across the cohort rather than restricted to a single molecular subtype.

Sequential integration of RNA-seq, poly(A)-site and CAGE evidence, followed by removal of reverse-transcription-switching artefacts, refined 129,335 initial transcript models to a curated catalogue of 90,929 variants, representing 70.3% of the initial set. Supported transcripts were retained across all four principal categories, with substantial numbers of NIC and NNC models remaining after curation **(Fig. 1c)**. The expanded non-reference repertoire was therefore not eliminated by orthogonal evidence filtering.

The curated NIC and NNC transcripts retained predominantly multi-exonic architectures and showed broader exon-count distributions than FSM and ISM transcripts **(Fig. 1d)**. Transcript diversity was also concentrated in a subset of genes, including COL1A1, FN1, COL3A1 and ERBB2, with the sample-group composition of detected isoforms varying among genes. ERBB2 isoforms were detected predominantly in HER2 samples, consistent with the expected enrichment of ERBB2 expression in this subtype (**Fig. 1e**). Alternative splicing event classes further showed distinct functional associations. Alternative first-exon events displayed the broadest enrichment profile, whereas retained introns and skipped exons were associated with multiple signalling, cell-cycle, immune and metabolic programmes; the remaining event classes showed more restricted enrichment patterns **(Fig. 1f)**. Extensive transcript diversity therefore persisted after stringent curation and showed non-random organization across genes and splicing-event classes. We next assessed how extensively this diversity propagated into translation-supported proteoform space.

### Alternative transcript diversity propagates extensively into translation-supported proteoform space

To determine how much of the curated transcript repertoire reached the translational level, we analysed publicly available Ribo-seq datasets from normal breast epithelial, ER-positive and triple-negative breast cancer cell lines. Ribosome-protected fragments were processed to identify P-site-supported candidate open reading frames and quantify their ribosome occupancy. Candidate ORF sequences were characterised by BLASTP searches against UniProt and HMMER searches against Pfam profiles. All detected candidate ORFs were initially retained to characterise gene-level ORF diversity and ORF-class composition. One primary Ribo-seq-supported ORF was subsequently selected for each transcript to establish a one-to-one correspondence between transcript isoforms and representative translation-supported ORFs for downstream proteoform analyses (**Fig. 2a**).

**Figure 2.**
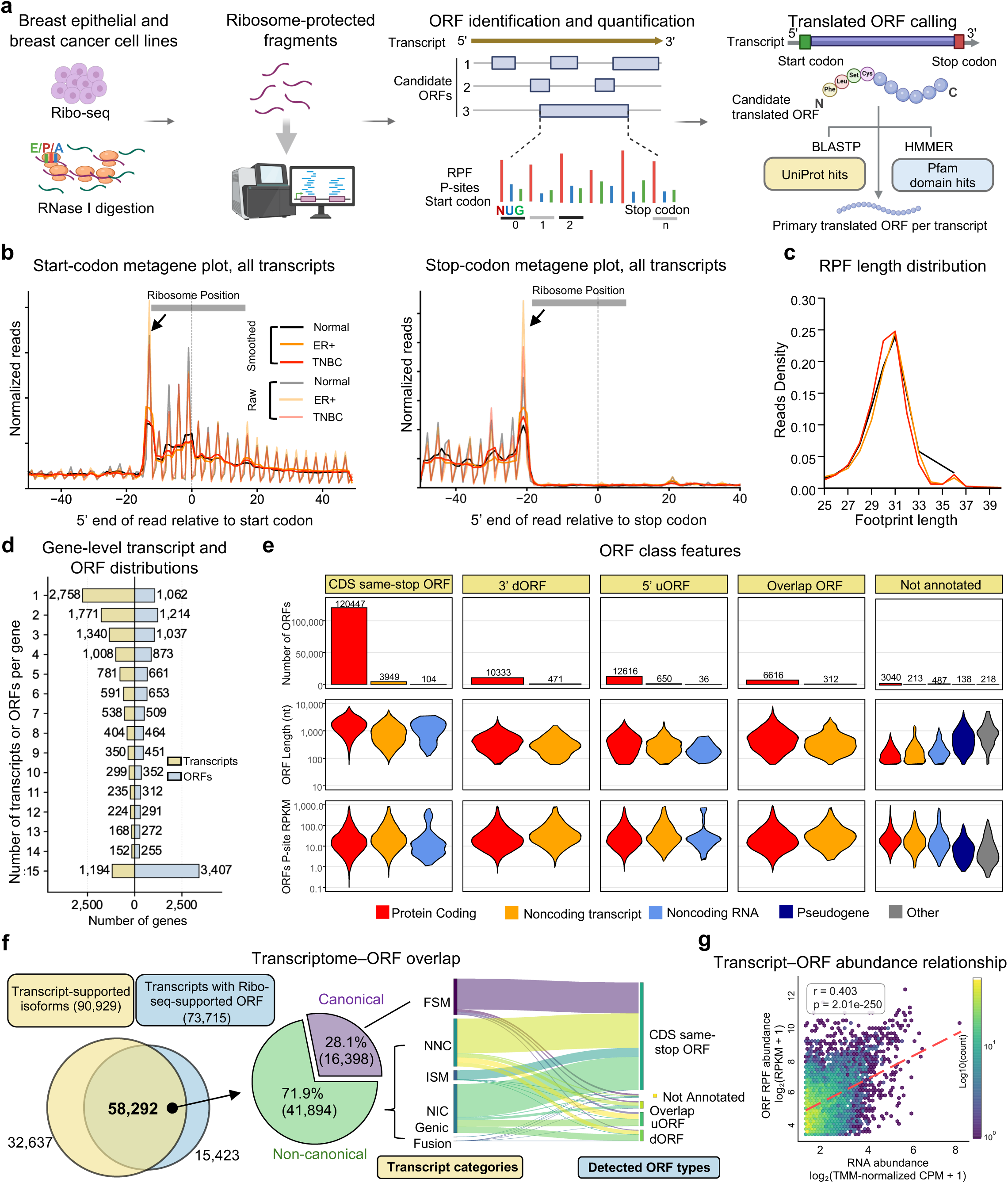
Ribosome profiling delineates translated open reading frames across breast epithelial and breast cancer cell lines. **a**, Workflow for processing publicly available Ribo-seq datasets, including RPF processing, ORF identification and quantification, protein sequence and domain annotation, and selection of a primary Ribo-seq-supported ORF for each transcript. **b,** Metagene profiles of normalized RPF reads around translation start and stop codons in normal, ER-positive (ER+) and triple-negative breast cancer (TNBC) cell lines; thin and thick lines denote raw and smoothed profiles, respectively. **c,** RPF length distributions across the three cell-line groups. **d,** Gene-level distributions of detected transcript isoforms and candidate ORFs. **e,** Numbers, lengths and RPF abundances of major candidate ORF classes, stratified by transcript biotype. **f,** Overlap between transcript-supported isoforms and transcripts assigned a primary Ribo-seq-supported ORF, together with the distribution of reference-matched and non-FSM transcript structures and their relationships with detected ORF classes. Transcript structural categories include full-splice match (FSM), incomplete-splice match (ISM), novel in catalog (NIC), novel not in catalog (NNC), genic and fusion transcripts. **g,** Relationship between transcript-level RNA abundance, quantified as log₂(TMM-normalized CPM + 1), and ORF-level RPF abundance, quantified as log₂(RPKM + 1). Pearson’s correlation coefficient and the two-sided *P* value are shown; colour indicates point density. RPF, ribosome-protected fragment.

The Ribo-seq datasets showed the positional and fragment-length characteristics expected of ribosome-protected reads. Aggregated RPF 5′-end profiles displayed enrichment at characteristic positions surrounding translation start and stop codons in all three cell-line groups (**Fig. 2b**). Most ribosome-protected fragments were 29-31 nucleotides long, with similar major peaks in normal, ER-positive and triple-negative breast cancer cells (**Fig. 2c**). These features supported transcript-resolved ORF identification across the combined datasets.

Analysis of all detected candidate ORFs revealed gene-level ORF diversity beyond transcript multiplicity alone. Gene-level distributions differed between reconstructed transcripts and candidate ORFs, with transcript isoforms enriched in lower-count categories and candidate ORFs showing a pronounced shift towards higher per-gene multiplicity. This difference was most evident in the ≥15 category, which contained 3,407 genes for candidate ORFs compared with 1,194 genes for transcript isoforms (**Fig. 2d**). CDS same-stop ORFs formed the largest candidate class and arose predominantly from protein-coding transcripts. RPF signals were also detected in 5′ upstream ORFs, 3′ downstream ORFs, overlapping ORFs and previously unannotated ORFs, including ORFs associated with noncoding transcript biotypes (**Fig. 2e**). These alternative ORF classes were generally shorter than CDS same-stop ORFs but retained measurable ribosome occupancy, showing that translation signals extended beyond conventional annotated coding regions.

Of the 90,929 high-confidence transcript isoforms identified from breast tissue, 58,292 were also represented among transcripts assigned a primary Ribo-seq-supported ORF, corresponding to 64.1% of the curated transcript repertoire under the assayed cell-line conditions (**Fig. 2f**). The overlapping set comprised 16,398 canonical and 41,894 non-canonical transcripts, with non-canonical transcript structures accounting for 71.9%. FSM, ISM, NIC, NNC, genic and fusion transcripts contributed to multiple ORF classes rather than mapping exclusively to a single coding architecture. Because the tissue transcript catalogue and Ribo-seq datasets were derived from different biological materials, the non-overlapping fractions should not be interpreted as evidence of absent translation in breast tissues or cell-line-specific translation.

Transcript-level RNA abundance was moderately correlated with ORF-level RPF abundance (Pearson’s r = 0.403, two-sided P = 2.01 × 10⁻²⁵⁰; **Fig. 2g**), indicating that variation in ribosome occupancy was not explained by transcript abundance alone. Ribo-seq therefore provided translational evidence beyond transcript abundance. Under the assayed cell-line conditions, nearly two-thirds of the curated transcript repertoire was linked to a primary Ribo-seq-supported ORF, and more than two-thirds of this overlapping set arose from non-canonical transcript structures. Alternative transcript diversity was therefore not confined to the RNA level.

### Translation-supported proteoforms exhibit coherent structural and cellular organization while retaining isoform-specific remodelling

We examined the full set of 73,715 primary Ribo-seq-supported ORFs, comprising 58,292 ORFs matched to the curated tissue transcript repertoire and 15,423 additional ORFs represented in the Ribo-seq transcript set. AlphaFold modelling yielded 53,066 protein structure models, of which 31,693 met a mean pLDDT threshold of at least 70. These high-confidence isoform-derived structures were combined with 14,908 representative structures from the human proteome to generate an integrated 3DisoGalaxy structure set comprising 46,601 proteins (**Fig. 3a**).

**Figure 3.**
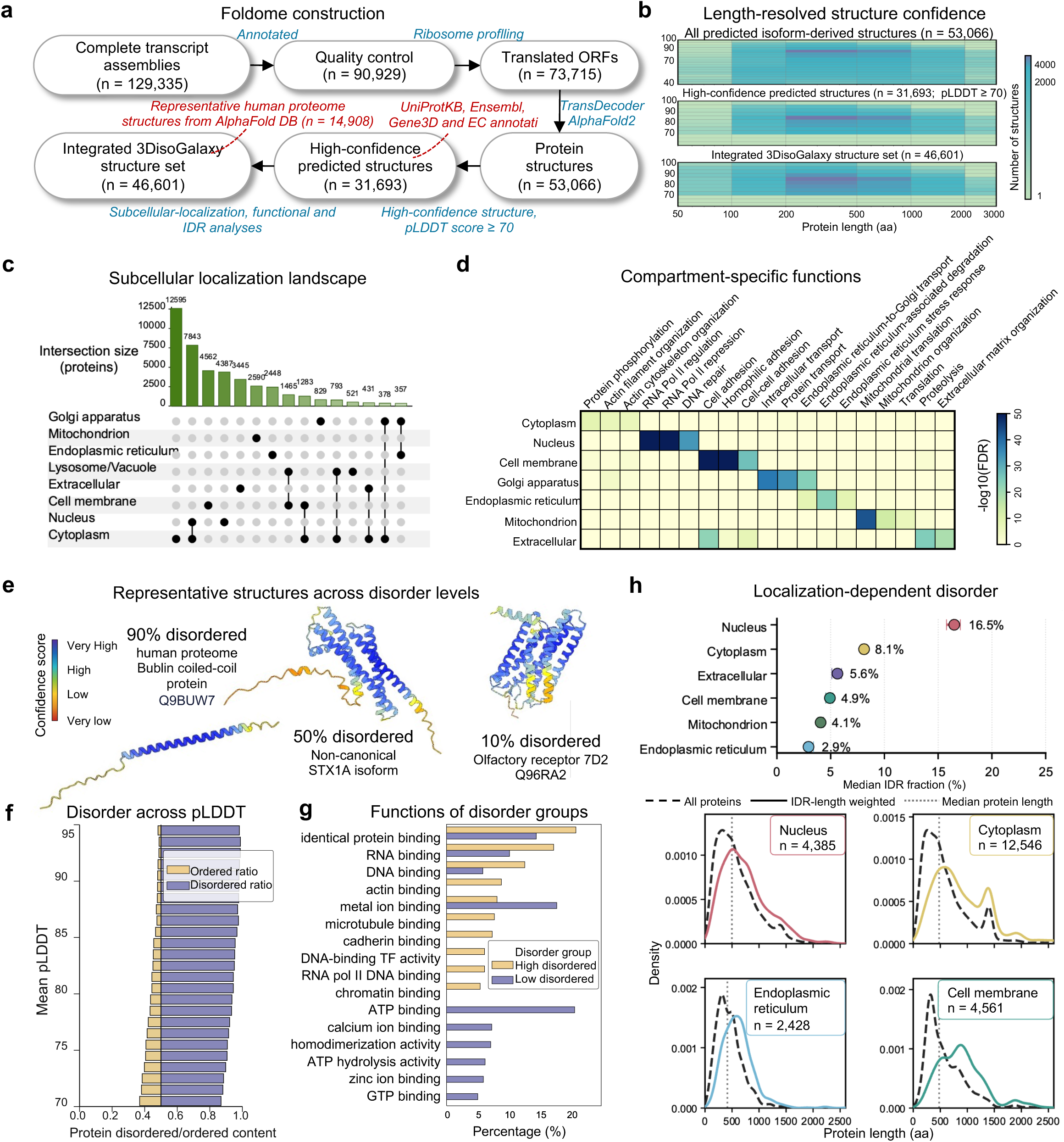
Structural and subcellular organization of the breast cancer foldome. **a**, Workflow integrating complete transcript assemblies (n = 129,335), quality-controlled transcripts (n = 90,929), translation-supported open reading frames (ORFs; n = 73,715) and AlphaFold-predicted protein structures (n = 53,066). Structures with pLDDT ≥ 70 (n = 31,693) were combined with 14,908 representative human proteome structures to construct a protein galaxy comprising 46,601 proteins. **b,** Length-resolved pLDDT density maps for all predicted structures, high-confidence structures and the combined protein galaxy. **c,** UpSet analysis showing the abundance and overlap of proteins assigned to major subcellular compartments. **d,** Enrichment of molecular functions across subcellular compartments. Colour represents −log10(FDR). **e,** Representative AlphaFold structures of proteins with high, intermediate and low predicted disorder content, coloured by structural confidence. **f,** Relative disordered and ordered sequence content across pLDDT intervals. **g,** Molecular-function distributions of proteins with high or low disorder content. **h,** Median intrinsically disordered region (IDR) fractions across subcellular compartments (top) and corresponding protein-length distributions for nuclear, cytoplasmic, endoplasmic reticulum and cell-membrane proteins (bottom). Dashed black curves represent all proteins, coloured curves represent IDR-length-weighted distributions and vertical dotted lines indicate median protein length.

To obtain orthogonal experimental support for representative non-canonical transcript models encoded within the final structure set, we selected HLA-C, SLC4A2 and SLC39A1 for RT-PCR analysis across normal mammary epithelial and breast cancer cell lines. Products of the expected sizes were detected for the corresponding canonical and candidate non-canonical transcript models (**Supplementary Fig. 3**). Sanger sequencing directly confirmed the predicted novel exon-exon junction in SLC4A2 and confirmed the identities of the HLA-C and SLC39A1 amplicons, although the latter two reads did not independently resolve transcript-specific splice junctions. These experiments provide targeted RNA-level support for representative transcript models selected from the structure atlas rather than experimental validation of the complete proteoform repertoire.

The predicted structures covered a broad range of protein lengths and structural-confidence values. High-confidence models were retained across the length spectrum, including both short proteins and proteins containing several thousand residues, rather than being restricted to a narrow protein-size interval (**Fig. 3b**). Confidence filtering therefore preserved structurally supported proteins across diverse sequence lengths.

Predicted subcellular localization and functional annotation showed concordant compartment-specific organization. Cytoplasmic and nuclear proteins formed the largest individual groups, while the observed intersections included proteins assigned to more than one cellular compartment (**Fig. 3c**). Nuclear proteins were associated with transcriptional regulation and DNA repair, cell-membrane proteins with cell adhesion, mitochondrial proteins with mitochondrial translation and organization, and extracellular proteins with proteolysis and extracellular-matrix processes. Cytoplasmic proteins were linked to protein phosphorylation and actin-associated processes, whereas Golgi-apparatus and endoplasmic-reticulum proteins were associated with intracellular transport, secretory-pathway trafficking, endoplasmic-reticulum-associated degradation and stress responses (**Fig. 3d**). The correspondence between predicted localization and compartment-characteristic functions indicated that the proteoform set retained expected relationships between cellular location and biological role.

Intrinsic disorder provided an independent measure of sequence and structural organization. Representative proteins containing approximately 90%, 50% and 10% predicted disorder showed a progression from extended regions with low structural confidence to compact structures containing continuous high-confidence cores (**Fig. 3e**). Across the analysed proteoforms, ordered sequence content increased with mean pLDDT, whereas disordered sequence was more prominent among lower-confidence models (**Fig. 3f**). Sequence-based disorder prediction and AlphaFold confidence thus captured related properties using independently derived measurements.

Molecular-function profiles also differed according to disorder content. Highly disordered proteins were more frequently associated with identical-protein binding, RNA and DNA binding, actin binding, transcription-factor activity and chromatin binding. Proteins with low disorder content showed greater representation of metal-ion, ATP, calcium-ion, zinc-ion and GTP binding, together with ATP hydrolysis and homodimerization activities, which are commonly mediated by compact folded domains (**Fig. 3g**). Disorder was also distributed unevenly across cellular compartments. Nuclear proteins had the highest median IDR fraction at 16.5%, followed by cytoplasmic proteins at 8.1%, extracellular proteins at 5.6%, cell-membrane proteins at 4.9%, mitochondrial proteins at 4.1% and endoplasmic-reticulum proteins at 2.9% (**Fig. 3h**). Disordered sequence therefore contributed to protein organization to different extents across cellular compartments.

We then assessed how isoform variation modified this broadly organized protein landscape. Isoform-reference comparisons encompassed N- and C-terminal truncations as well as internal insertions and deletions, each of which could modify protein architecture (**Supplementary Fig. 4a**). Among the 46,411 isoforms represented in this analysis, 38,903 (83.8%) originated from multi-isoform genes, whereas 7,508 (16.2%) originated from singleton genes (**Supplementary Fig. 4b**). Most of the analysed isoform space therefore arose from genes producing multiple protein forms.

Pfam architecture was retained in a large proportion of isoform-reference comparisons. Proteoforms with unchanged and remodelled Pfam composition were observed across a broad range of pLDDT values, and the distribution of changes in unique Pfam count was strongly centred at zero (**Supplementary Fig. 4c,d**). A subset of proteoforms nevertheless showed domain gains, domain losses or altered domain combinations. Because these events were not confined to proteins with low structural confidence, Pfam remodelling was unlikely to arise solely from poorly modelled sequences. Alternative splicing therefore commonly retained the principal domain architecture while modifying selected structural modules in individual isoforms.

Isoform variation also affected sequence features outside annotated domains. Changes in IDR length and IDR fraction occurred in both positive and negative directions, indicating that alternative isoforms could gain or lose disordered sequence relative to their reference proteins (**Supplementary Fig. 4e**). Predicted localization showed a similarly heterogeneous pattern, with isoform families differing in the extent of localization gain, loss and reconfiguration (**Supplementary Fig. 4f**). Among isoform-rich cell-membrane genes, localization remodelling varied substantially and did not consistently track the extent of Pfam alteration (**Supplementary Fig. 4g**). Changes in predicted cellular targeting could therefore occur without extensive restructuring of annotated protein domains.

Translation-supported proteoforms retained recognizable structural and cellular organization, but this organization did not imply that related isoforms were equivalent. The principal protein framework was frequently preserved while individual domains, disordered regions and localization-associated features were modified. We therefore tested whether short regulatory motifs represented an additional layer of localized proteoform remodelling (**Fig. 4**).

**Figure 4.**
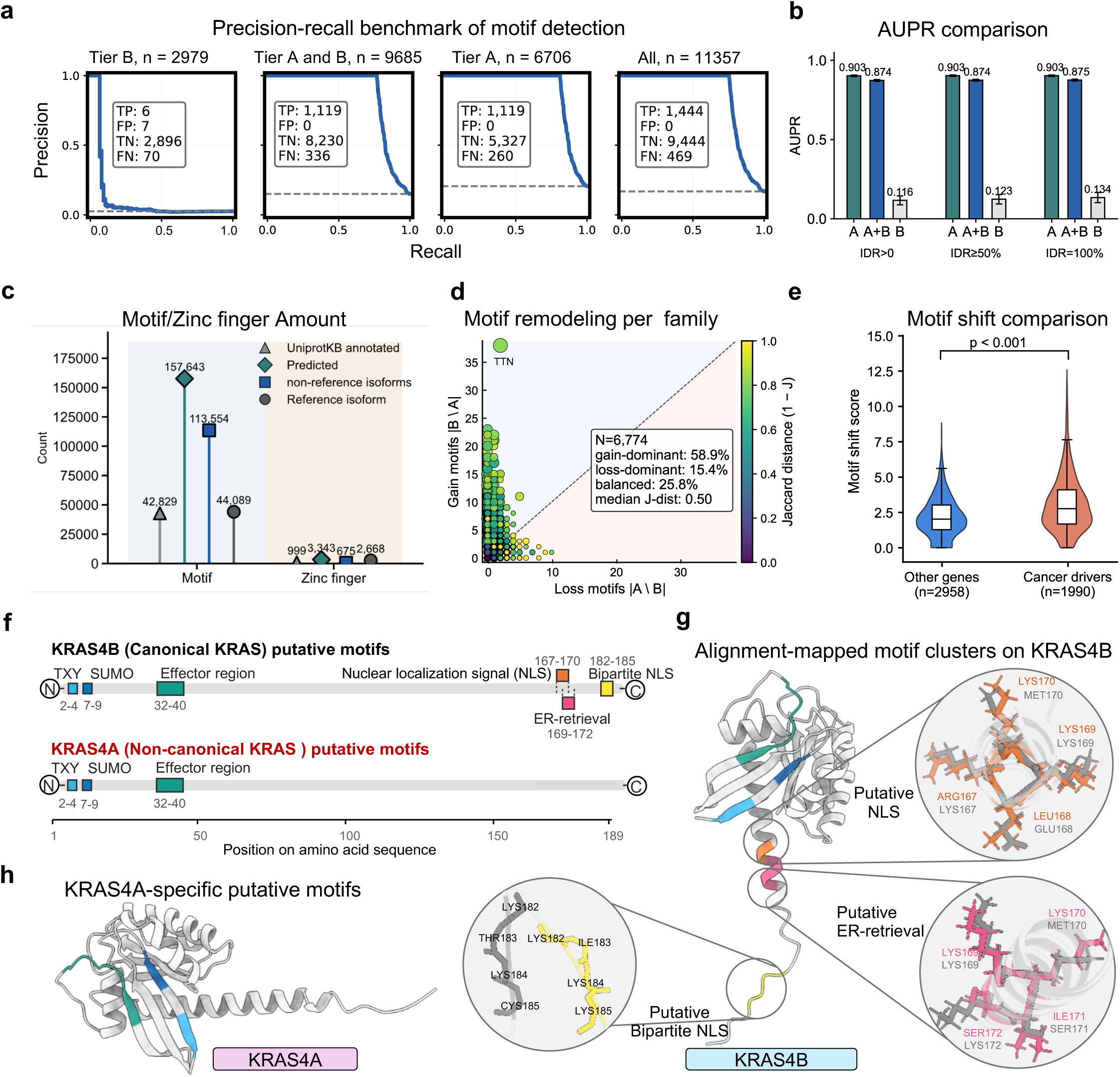
Benchmarking and isoform-level analysis of motif remodelling. **a**, Prediction-level precision–recall ranking of retrieved structural motif matches across Tier B, combined Tier A and B, Tier A and complete benchmark sets. *n* denotes the number of benchmark instances. **b,** Comparison of area under the precision–recall curve (AUPR) for Tier A, combined Tier A and B, and Tier B motif matches under the indicated IDR thresholds (IDR > 0, IDR ≥ 50% and IDR = 100%). **c,** Numbers of UniProtKB-annotated and putative structure-supported motifs and zinc-finger elements in reference and non-reference isoforms. **d,** Numbers of motif classes unique to the reference repertoire (A \ B) or pooled non-reference repertoire (B \ A) across gene-level isoform families. Points are coloured by the Jaccard distance between the reference and pooled non-reference motif repertoires; the diagonal indicates equal numbers of motif classes unique to each repertoire. The inset summarizes the proportions of gain-dominant, loss-dominant and balanced families among the 6,774 gene-level isoform families analysed. **e,** Distribution of motif-shift scores in cancer-driver genes (*n* = 199) and other genes (*n* = 2,998). Statistical significance was assessed using a two-sided Wilcoxon rank-sum test. Boxes show the median and interquartile range, whiskers extend to 1.5 times the interquartile range, and violin widths represent kernel-density estimates. **f,** Linear maps of putative structure-supported motifs in the reference KRAS4B isoform and the alternative KRAS4A isoform, showing shared N-terminal motifs and isoform-specific motifs within the C-terminal hypervariable region. **g,** Alignment-mapped motif clusters visualized on the predicted KRAS4B structural model, highlighting putative NLS-like and other basic motifs within the C-terminal hypervariable region. **h,** Comparison of the predicted KRAS4A and KRAS4B structural models, showing the mapped locations of isoform-specific putative C-terminal motifs, including the putative bipartite NLS-like motif. NLS, nuclear localization signal.

### Localized motif remodelling differentiates proteoforms and is elevated in cancer-driver genes

The structural coherence of translation-supported proteoforms did not imply regulatory equivalence. We examined whether related proteoforms differed through the gain, loss or repositioning of short regulatory motifs. To obtain a conservative structure-informed motif set, Folddisco was used to search protein structures for consensus-matching structural motifs under stringent geometric and coverage criteria (Kim et al., 2026). Performance was evaluated using tiered benchmark sets representing different levels of structural-match stringency.

Among retrieved candidate regions, high-stringency Tier A matches achieved a prediction-level average precision of 0.903 for overlap with curated UniProt motif-bearing regions. The combined Tier A and B set yielded prediction-level average precision values of 0.874-0.875 across the tested intrinsically disordered region (IDR) thresholds (**Fig. 4a,b**; **Supplementary Fig. 5a-d**). Performance of the higher-stringency match sets remained largely stable across increasingly restrictive IDR thresholds. Moderate-stringency Tier B matches showed substantially lower prediction-level average precision, ranging from 0.116 to 0.134 across the tested thresholds. Decoy-based controls yielded low estimated false-discovery rates at the stringent operating point (**Supplementary Fig. 5d**). Tier A matches satisfying the IDR ≥ 50% criterion were retained as the conservative default for downstream isoform-level comparisons.

Application of this framework expanded regulatory annotation across the proteoform repertoire. In addition to 42,829 UniProtKB-annotated motifs, 157,643 putative motifs were identified computationally, comprising 44,089 assignments in reference isoforms and 113,554 in non-reference isoforms (**Fig. 4c**). A further 3,343 structure-supported zinc-finger assignments were detected, including 2,668 in reference isoforms and 675 in non-reference isoforms. These predictions extended motif annotation beyond reference proteins and enabled matched comparisons of regulatory sequence features within proteoform families.

Motif remodelling was quantified across 6,774 gene-level isoform families by comparing the pooled motif repertoire of the reference proteoform with the union of motifs detected across its corresponding non-reference proteoforms. Motif gains were defined as motif classes present only in the pooled non-reference repertoire, whereas losses were defined as motif classes present only in the reference repertoire. Differences generally arose through discrete motif gains and losses rather than complete replacement of the motif repertoire (**Fig. 4d**). In this gene-level pooled comparison, 58.9% of families were gain-dominant, 15.4% were loss-dominant and 25.8% were balanced. Gain counts exceeded loss counts in the pooled gene-level comparison (Wilcoxon signed-rank test, P < 0.001), although this asymmetry should be interpreted in the context of the larger number of non-reference isoforms contributing to the pooled repertoire. Representative gain-dominant and loss-dominant proteoform families are shown in **Supplementary Fig. 6**.

Cancer-driver families showed higher motif-shift scores than the remaining gene set (two-sided Wilcoxon rank-sum test, P < 0.001; **Fig. 4e**). The distributions overlapped, indicating that motif remodelling was not exclusive to cancer-associated genes. The shift towards higher scores indicated greater motif-repertoire divergence among cancer-driver gene families than among other genes.

KRAS provided an example of regulatory remodelling within a preserved structural framework. KRAS4A and KRAS4B shared the central Ras fold, the Ras effector-associated region and N-terminal TXY- and SUMO-associated motif annotations (**Fig. 4f-h**). Their divergent C-terminal hypervariable sequences, however, produced distinct candidate motif organizations. KRAS4B contained additional predicted C-terminal elements, including NLS-like, ER-retrieval-like and bipartite NLS-like motifs that were not detected in the corresponding KRAS4A sequence (**Fig. 4f**). Structural mapping placed these motif clusters within the flexible KRAS4B C-terminal region without disrupting the conserved Ras core (**Fig. 4g,h**). The KRAS comparison illustrated how alternative terminal sequences can modify candidate localization and interaction determinants while preserving the principal folded domain.

Localized motif-repertoire remodelling was widespread across gene-level isoform families and was more pronounced in cancer-driver genes. We used the integrated structure network to identify cancer-associated candidates that retained a recognizable protein core while showing disruption of a major regulatory module.

### 3DisoGalaxy places isoform-derived proteins within established fold space and prioritizes an AKT1-ΔPH candidate

We constructed a structure-similarity network from the 46,601 proteins represented in 3DisoGalaxy, connecting protein pairs with a structure similarity cutoff (TM-score of at least 0.9) **(Fig. 5a)**. Structurally connected proteins generally showed concordant Pfam annotations, although partially concordant domain compositions were more frequent in the largest network components **(Supplementary Fig. 7a,b)**.

**Figure 5.**
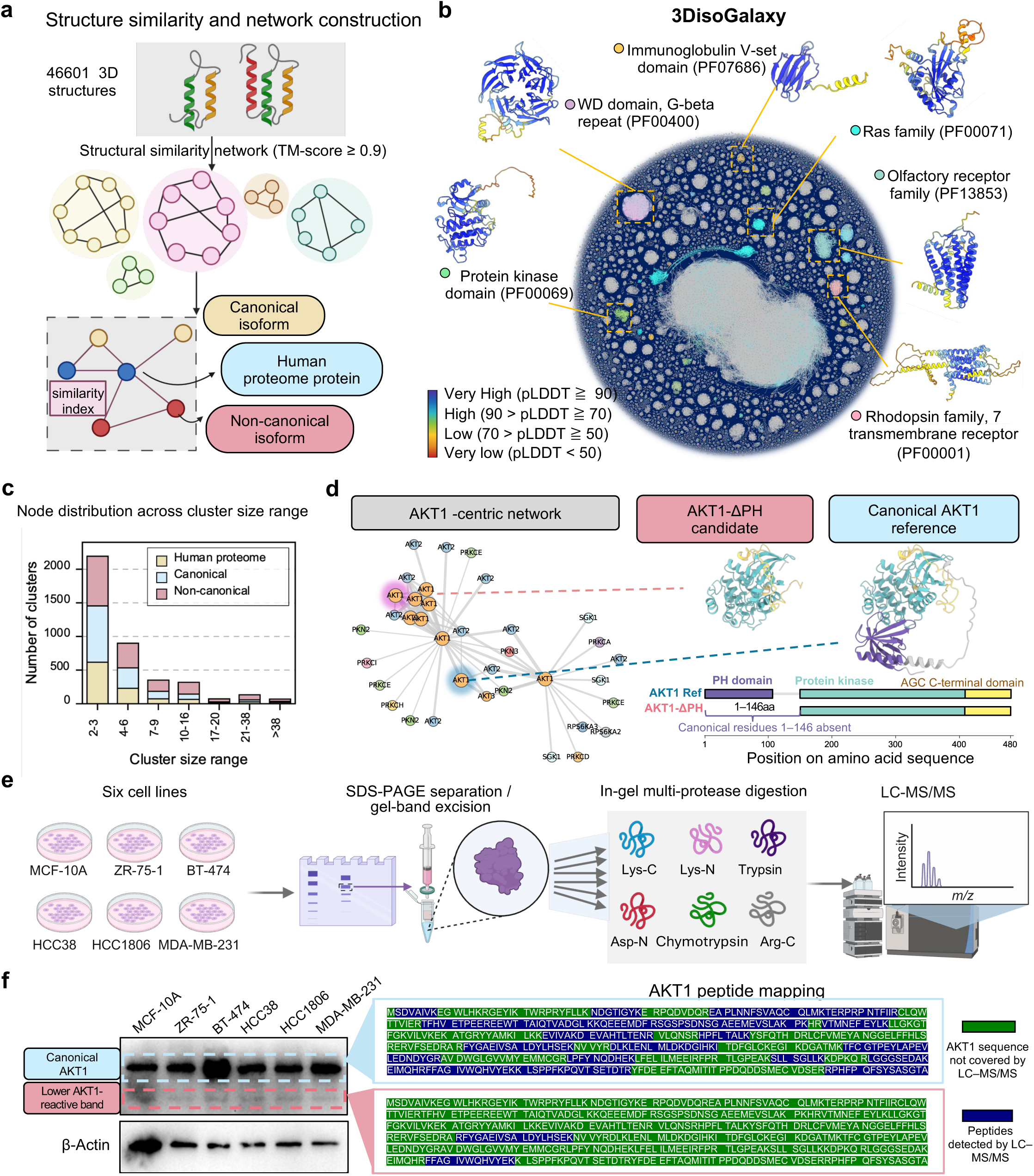
Structural organization of reference and isoform-derived proteins in 3DisoGalaxy and protein-level evidence consistent with a shorter AKT1 candidate. **a**, Construction of a structure-similarity network from 46,601 protein structures, comprising representative human proteome proteins, canonical isoforms and non-canonical isoforms, with edges defined by a TM-score of ≥0.9. **b,** Global 3DisoGalaxy structure-similarity network with representative Pfam families highlighted. **c,** Composition of non-singleton similarity-defined network components across component-size ranges and protein categories. **d,** AKT1-centred subnetwork and structural comparison of canonical AKT1 with the predicted 334-aa AKT1-ΔPH candidate, which lacks canonical residues 1–146 and the PH domain while retaining the protein kinase and AGC-kinase C-terminal domains. **e,** Proteomic workflow involving gel-band excision, parallel multi-protease digestion and LC–MS/MS. **f,** Immunoblot detection of canonical-size and lower-molecular-weight AKT1-reactive bands, with β-actin as a loading control. AKT1-derived peptides identified from the corresponding gel regions were mapped to the canonical AKT1 sequence; blue indicates detected peptide coverage and green indicates positions without coverage.

The resulting network recovered established protein architectures, including protein kinases, Ras-family proteins, WD-repeat proteins, immunoglobulin V-set domains, olfactory receptors and rhodopsin-family seven-transmembrane receptors **(Fig. 5b)**. Canonical isoforms, non-canonical isoforms and representative human proteome proteins occupied shared structural neighbourhoods. Non-canonical isoforms were distributed across the full range of non-singleton component sizes, including the largest network components, indicating that they were not restricted to small or structurally isolated regions of the network **(Fig. 5c)**.

We next focused on a non-canonical AKT1 isoform, hereafter termed the AKT1-ΔPH candidate. Although grouped within the AKT1-centred neighbourhood, this candidate was positioned apart from canonical AKT1 and the other AKT1 isoforms, indicating substantial structural divergence within the isoform family (**Fig. 5d**). Consistent with its distinct network position, 3DisoDeepPF predicted that the pleckstrin homology domain present in the other AKT1 isoforms was absent from AKT1-ΔPH, whereas the kinase-associated domains were retained (Jiang et al., 2026). The convergence of network-level structural divergence and predicted PH-domain loss led us to prioritize AKT1-ΔPH for further structural and experimental investigation **(Fig. 5d)**.

Immunoblotting revealed a lower-molecular-weight AKT1-reactive band in addition to canonical-size AKT1, and LC–MS/MS identified AKT1-derived peptides in the corresponding gel region **(Fig. 5e,f)**. Because the detected peptides mapped to regions shared with canonical AKT1, they support the presence of a shorter AKT1 species but do not resolve its complete sequence or precise N-terminal boundary. AKT1-ΔPH therefore provides protein-level support for the central pattern identified across 3DisoGalaxy: preservation of a recognizable functional core alongside selective loss of a regulatory module. Its retained kinase architecture is consistent with potential AKT1 catalytic capacity, whereas loss of the PH domain predicts altered canonical membrane recruitment and regulatory control. This case demonstrates how 3DisoGalaxy can progress from proteoform-scale organization to a biologically interpretable candidate for direct functional investigation.

## Discussion

The extent to which full-length alternative transcript diversity materialises as protein diversity remains unresolved. Transcript detection alone does not establish productive translation, ribosome engagement does not necessarily imply stable protein accumulation, and conventional proteomics has limited sensitivity for distinguishing closely related protein isoforms (Ezkurdia et al., 2015; Liu et al., 2017; Lau et al., 2019). By integrating full-length transcript reconstruction, ribosome profiling and structure–function inference, this study addresses three successive questions: how extensively alternative transcripts enter translation-supported sequence space, whether the resulting proteoforms exhibit coherent protein-level organisation, and how related proteoforms acquire distinct regulatory properties. We show that alternative transcript diversity is widely represented among translation-supported ORFs, that the encoded products conform to established relationships among protein structure, intrinsic disorder, localisation and function, and that proteoform divergence is frequently concentrated in modular regulatory features rather than requiring the generation of entirely new protein folds. The structure–function atlas 3DisoGalaxy further enabled the identification of an AKT1-ΔPH proteoform that retains the kinase core while lacking an intact pleckstrin homology domain, providing an experimentally supported example of regulatory remodelling imposed on a preserved functional core.

A central observation was that alternative transcript diversity was not largely confined to the RNA level. Of the 90,929 curated full-length transcripts, 58,292, or 64.1%, were represented among transcripts assigned a primary Ribo-seq-supported ORF under the analysed cell-line conditions. Non-canonical transcript structures accounted for 71.9% of this overlapping set. These proportions indicate that translation-supported sequence diversity is broader than would be inferred from the relatively small number of alternative protein isoforms resolved through isoform-discriminating peptides. Closely related proteoforms share most of their amino acid sequences, leaving few unique peptides, and these peptides may not be generated, recovered or detected efficiently in conventional bottom-up proteomic workflows (Ezkurdia et al., 2015; Liu et al., 2017; Lau et al., 2019). Transcript-informed proteomic searches and intact-protein approaches have identified additional non-canonical protein forms, indicating that technical observability contributes to the apparent gap between transcript and protein diversity (Tran et al., 2011; Lau et al., 2019).

Ribosome profiling provides an independent measure of ribosome occupancy and enables translated ORFs to be identified at high positional resolution (Ingolia et al., 2009; Ingolia et al., 2019). It does not, however, establish that every ribosome-associated sequence accumulates as a stable, abundant or functional protein. The tissue transcriptomes and cell-line Ribo-seq datasets analysed here were also not sample matched. The observed 64.1% overlap should therefore not be interpreted as a direct estimate of transcript translation within individual breast tumours, and the non-overlapping tissue transcripts should not be considered untranslated. Rather, the results establish that a substantial fraction of the curated transcript repertoire is represented in translation-supported ORF space across relevant breast epithelial and cancer contexts. This conclusion is consistent with isoform-resolved ribosome-profiling studies showing that alternative transcript usage can propagate into differential translation (Reixachs-Solé et al., 2020).

Translation support alone is insufficient to distinguish biologically organised protein products from incidental or non-productive translation. A second layer of evidence therefore came from the structural and cellular properties of the encoded sequences. The translation-supported proteoforms spanned a broad range of protein lengths and structural-confidence values, and high-confidence filtering did not selectively retain only short or structurally simple proteins. Across the resulting foldome, protein length, AlphaFold confidence and intrinsic disorder followed expected relationships. Regions with low AlphaFold confidence frequently correspond to intrinsically disordered or context-dependent sequences, although low confidence should not be treated as equivalent to disorder in every case (Tunyasuvunakool et al., 2021; Akdel et al., 2022). Highly disordered proteins were enriched for nucleic-acid binding, transcriptional regulation and interaction-related functions, consistent with the established roles of disordered regions in molecular recognition, signalling and regulatory control (Wright and Dyson, 2015; Holehouse and Kragelund, 2024).

Predicted localisation provided an independent cellular context for this organisation. Nuclear proteins contained the highest median proportion of disordered sequence, whereas proteins assigned to membranes, mitochondria and the secretory pathway were generally more ordered. Compartment assignments were accompanied by corresponding functional programmes, including transcriptional and DNA-associated processes in the nucleus, adhesion at the cell membrane, trafficking through the Golgi apparatus and endoplasmic reticulum, mitochondrial translation and extracellular-matrix organisation. These relationships are consistent with experimentally informed maps of the spatial organisation of the human proteome (Thul et al., 2017). Their convergence across independently derived structure, disorder, localisation and functional annotations supports the biological coherence of the repertoire as a whole, although it does not demonstrate the stability or function of every individual predicted proteoform.

Structural similarity analysis extended this interpretation by placing non-canonical proteoforms within established human fold space. Alternative proteoforms occurred across both small and large structural clusters and were represented in recognisable kinase, Ras-family, WD-repeat, immunoglobulin and membrane-receptor neighbourhoods. AlphaFold has substantially expanded structural coverage of the human proteome, while its predicted structures remain hypotheses that require appropriate confidence assessment and experimental validation for specific biological applications (Jumper et al., 2021; Tunyasuvunakool et al., 2021; Akdel et al., 2022). Within these limits, our findings indicate that non-canonical proteoforms are not generally restricted to an isolated set of structurally anomalous sequences. Much of the translation-supported repertoire is compatible with established fold families, suggesting that alternative splicing commonly modifies proteins within the constraints of existing structural architectures.

The combined analyses support a hierarchy of proteoform diversification. Major folded domains and catalytic cores were often retained, whereas alternative sequence usage more frequently modified terminal regions, intrinsically disordered segments, localisation determinants and short regulatory motifs. Previous studies have similarly shown that alternatively spliced regions are enriched in disordered sequences, interaction motifs and post-translational modification sites, and that these changes can remodel protein-interaction networks (Buljan et al., 2012; Ellis et al., 2012; Yang et al., 2016; Lau et al., 2019). Domain gain or loss can introduce or remove a discrete structural module, while smaller changes in flexible regions may alter accessibility, interaction specificity or cellular targeting without disrupting the principal fold. Intrinsically disordered regions are particularly suited to this form of regulation because they can accommodate multiple binding motifs and modification sites within conformationally flexible sequences (Wright and Dyson, 2015; Holehouse and Kragelund, 2024).

Short regulatory motifs provide an especially plastic layer of proteoform diversification. Their limited sequence requirements allow motifs to be gained, lost or repositioned through relatively small sequence changes, potentially affecting partner binding, localisation, degradation or signalling (Van Roey et al., 2014). Among the 6,774 isoform families examined here, motif remodelling was predominantly gain-biased, with 58.9% of families classified as gain-dominant compared with 15.4% classified as loss-dominant. Cancer-driver genes also showed higher motif-shift scores than other genes. This association does not establish that the predicted motif changes are oncogenic, but it is consistent with evidence that alternative splicing can expand isoform-specific interaction capabilities and contribute to cancer-associated functional changes (Yang et al., 2016; Kahles et al., 2018).

The comparison between KRAS4A and KRAS4B illustrates this regulatory principle. The two isoforms retain a common Ras GTPase fold but differ in their alternatively encoded C-terminal regions, which contain membrane-targeting and trafficking determinants. Experimental studies have shown that the KRAS splice variants differ in structural dynamics, thermal stability, effector interactions and responses to trafficking proteins despite retaining closely related catalytic cores (Whitley et al., 2024). The NLS-like and trafficking-associated motifs predicted here should therefore be considered testable hypotheses rather than established localisation signals. Their concentration in the divergent C-terminal region nevertheless illustrates how alternative terminal sequences can remodel candidate spatial and interaction determinants while preserving the principal Ras architecture.

AKT1-ΔPH provides a more direct example of how structure–function inference can move from global description to proteoform discovery. The predicted proteoform lacks the N-terminal sequence required to form an intact pleckstrin homology domain but retains the kinase domain and C-terminal hydrophobic motif. The AKT pleckstrin homology domain binds phosphoinositides and couples AKT to membrane recruitment, conformational activation and phosphorylation by upstream kinases (Carpten et al., 2007; Manning and Toker, 2017). Its disruption therefore predicts altered spatial regulation and upstream input coupling rather than simple deletion of the catalytic machinery. Retention of the kinase core raises the possibility that AKT1-ΔPH preserves catalytic potential while operating under a different regulatory regime, although this possibility requires direct experimental testing.

The lower-molecular-weight AKT1-reactive band and LC–MS/MS peptide evidence provide orthogonal protein-level support for the existence of a shorter AKT1 form. The detected peptides map to regions retained in the predicted proteoform, but because they are shared with canonical AKT1, they do not independently establish its complete sequence or precise N-terminal boundary. The evidence therefore supports a shorter AKT1 species consistent with the predicted AKT1-ΔPH architecture rather than definitively resolving the intact proteoform. This distinction is important because bottom-up proteomics identifies proteolytic peptides rather than complete molecular forms, whereas top-down proteomics is better suited to resolving intact protein isoforms and their molecular modifications (Tran et al., 2011; Smith and Kelleher, 2013).

The identification of AKT1-ΔPH also clarifies the role of computational proteoform biology. This approach is not intended to replace direct biochemical characterisation or intact-protein proteomics. Its value lies in integrating complementary evidence to identify transcript-derived protein sequences that are sufficiently supported and biologically interpretable to warrant experimental investigation. Full-length transcripts define sequence architecture, Ribo-seq provides evidence of ribosome engagement, structural models reveal preserved and altered modules, and disorder, localisation and motif analyses generate mechanistic hypotheses. This framework converts a large transcript catalogue into a set of sequence-defined and experimentally prioritised proteoform candidates.

Several limitations define the scope of the conclusions. The long-read tissue data and cell-line Ribo-seq datasets originated from different biological materials, preventing sample-level coupling of transcript expression and translation. Ribosome engagement cannot distinguish stable proteins from rapidly degraded or low-abundance products. AlphaFold confidence, disorder prediction, localisation assignment and motif matching provide inferential rather than direct functional evidence. Predictions involving flexible tails, membrane-associated conformations or interaction-dependent structures require particular caution because these states may not be represented accurately by isolated single-chain models (Tunyasuvunakool et al., 2021; Akdel et al., 2022). The present analysis also focuses primarily on sequence-defined proteoforms arising from transcript variation and does not capture the additional diversity produced by post-translational modifications, proteolytic processing or combinations of molecular states (Smith and Kelleher, 2013).

For AKT1-ΔPH, isoform-specific N-terminal peptides, targeted mass spectrometry, N-terminomics or top-down proteomics will be required to resolve the intact protein sequence. Functional studies should determine its localisation, phosphorylation state, catalytic activity, substrate selection and response to pathway inhibitors. More generally, matched tumour transcriptomics, Ribo-seq and proteomics will be needed to quantify proteoform production within individual disease contexts. Isoform-specific perturbation and protein-level assays will then be required to establish whether the predicted regulatory changes produce measurable signalling or therapeutic phenotypes.

Within these boundaries, full-length alternative transcript diversity extends extensively into a translation-supported and biologically organised protein repertoire. Much of this variation is accommodated within established fold space, while potential functional differences are encoded through modular remodelling of domains, disordered regions, localisation determinants and short regulatory motifs. The increased motif remodelling observed in cancer-driver genes further indicates that this regulatory plasticity is unevenly distributed across the proteome. By linking transcript discovery to translation, protein organisation and targeted validation, 3DisoGalaxy provides a route from isoform cataloguing to experimentally testable proteoform biology and identifies AKT1-ΔPH as a candidate example of a conserved kinase core embedded within an altered regulatory architecture.

## Methods

### Reference annotations and protein structures

GENCODE comprehensive gene annotation release 41 and the GRCh38 reference genome were used throughout the human transcript analyses (Frankish et al., 2023). MANE Select annotations were used to define representative reference transcripts where available (Morales et al., 2022). Reviewed human protein annotations and annotated isoform sequences were obtained from UniProtKB/Swiss-Prot (UniProt Consortium, 2023). Human AlphaFold Database models (UniProt proteome UP000005640, database version 4) were used as representative reference structures (Varadi et al., 2022).

### Long-read transcript discovery and curation

#### Cross-sample transcript integration

Full-length PacBio transcript models were integrated across samples with TALON v5.0 using GENCODE v41 as the reference annotation (Wyman et al., 2020). Long reads were aligned with minimap2 and processed with SAMtools to generate alignment files containing MD tags (Li, 2018; Danecek et al., 2021). Reads were added to the TALON database using minimum alignment coverage and identity thresholds of 0.95. Transcripts detected in at least two independent samples were carried forward to annotation and quality control.

#### Transcript classification and quality control

Transcript models were classified and quality-controlled with SQANTI3 using GENCODE v41, CAGE peaks, poly(A) motifs and curated poly(A)-site resources (Pardo-Palacios et al., 2024). Transcripts were assigned to full-splice-match (FSM), incomplete-splice-match (ISM), novel-in-catalogue (NIC), novel-not-in-catalogue (NNC), genic, intronic, antisense, fusion or intergenic categories according to their splice-junction and transcript-boundary relationships with the reference annotation.

Potential indel-derived artefacts were corrected by realignment to GRCh38 and reevaluation with SQANTI3. Transcripts were required to have long-read support in at least three Iso-Seq samples for inclusion in the final curated catalogue. Novel junctions supported by fewer than five short reads in STAR splice-junction output were removed. Transcript 3′ ends located more than 100 bp from an annotated termination site and lacking support from curated poly(A)-site resources were also excluded. Reverse-transcription-switching artefacts identified by SQANTI3 were removed. This workflow reduced 129,335 initial transcript models to a curated catalogue of 90,929 transcripts.

### Alternative-splicing analysis

Alternative-splicing events were generated from the TALON-filtered long-read GTF using SUPPA2 (Alamancos et al., 2015; Trincado et al., 2018). Isoform-level TPM estimates from RSEM were supplied as the expression input. The analysed event classes comprised skipped exons, mutually exclusive exons, retained introns, alternative 3′ splice sites, alternative 5′ splice sites and alternative first and last exons. Differential splicing between basal and non-basal sample groups was assessed with SUPPA2 diffSplice using the empirical method and the corresponding PSI and isoform-expression matrices. Analyses were performed separately for each event class.

### Ribo-seq processing and translation-supported ORF identification

#### Read preprocessing and alignment

Raw Ribo-seq and matched RNA-seq FASTQ files were assessed with FastQC, and library-level summaries were compiled with MultiQC (Ewels et al., 2016). Adapter sequences were removed with cutadapt v4, using a maximum error rate of 0.05 and a minimum retained read length of 20 nt (Martin, 2011). Reads of 20–35 nt were retained for initial Ribo-seq processing. Reads aligning to ribosomal RNA or the Illumina PhiX genome were removed with Bowtie v1.3 (Langmead et al., 2009). The remaining reads were aligned to GRCh38.p13 with STAR v2.7 using the merged GENCODE and long-read transcript annotation (Dobin et al., 2013). Alignments were restricted to a maximum of two mismatches and uniquely mapped reads (-- outFilterMultimapNmax 1) and were generated in both coordinate-sorted and transcriptome-coordinate formats. Alignment files were processed and indexed with SAMtools (Danecek et al., 2021).

Read-length distributions, mapping rates, replicate correlations, P-site frame periodicity and metagene profiles surrounding annotated start and stop codons were evaluated as Ribo-seq quality-control measures. P-site offsets were calibrated by read length using RiboCode metagene profiles, with the principal analysis restricted to footprints of 26–34 nt (Xiao et al., 2018).

#### ORF detection, quantification and representative-ORF selection

Translated ORFs were detected with RiboCode v1.2.13 using the calibrated P-site offsets and the merged transcript annotation (Xiao et al., 2018). RiboCode calls included annotated coding sequences, upstream ORFs, downstream ORFs, overlapping ORFs and previously unannotated ORFs. ORF-type labels describe the position of an ORF relative to the associated annotated coding sequence and were treated separately from the canonical or non-canonical classification of the full transcript.

ORFs were required to exhibit significant three-nucleotide periodicity, enrichment of footprints in the zero frame and at least 15 assigned P-site counts. ORF abundance was quantified with ORFcount using 15-nt and 5-nt start- and stop-proximal exclusions, respectively, a 100-nt exon extension, a footprint-length range of 24–35 nt and strict P-site counting. After removal of ORFs shorter than 100 amino acids and exact duplicate peptide sequences, the longest RiboCode-supported ORF was retained as the primary ORF for each transcript. This produced a one-to-one mapping between transcripts and primary translation-supported protein sequences for downstream proteoform analysis. The correspondence between the tissue-derived catalogue and the cell-line Ribo-seq catalogue was interpreted as translation support under the assayed cell-line conditions, rather than direct evidence of translation in the original tissue specimen.

### ORF prediction and protein annotation

Candidate coding sequences were independently predicted from full-length transcript sequences with TransDecoder v5.7.1. Long ORFs extracted with TransDecoder.LongOrfs were searched against a human UniProt reference proteome using BLASTP v2.12.0 (max_target_seqs = 1; E-value ≤ 1 × 10−5) and against Pfam-A v35.0 using HMMER3 v3.3.2 (hmmscan; E-value ≤ 1 × 10−10) (Camacho et al., 2009; Eddy, 2011; Mistry et al., 2021). Homology and Pfam-domain evidence were incorporated with TransDecoder.Predict to annotate plausible protein-coding sequences. For the structure atlas, both canonical and non-canonical transcript classes were eligible, provided that their representative ORFs passed the RiboCode-support, length and redundancy filters described above.

### Protein-structure modelling and foldome construction

#### AlphaFold modelling and structure quality control

Protein structures were predicted with AlphaFold2.3 through ColabFold v1.5.5 (Jumper et al., 2021; Mirdita et al., 2022). Multiple-sequence alignments were generated with colabfold_search against a local ColabFold database using environmental sequences and without structural templates. Structure prediction was performed with colabfold_batch, using three recycles and three models per sequence. The top-ranked model was subjected to one round of Amber relaxation. Predictions were configured to stop when the internal score reached 85. Models with a mean predicted local distance difference test (pLDDT) score of at least 70 were retained for downstream analysis, consistent with established interpretation of AlphaFold confidence scores (Tunyasuvunakool et al., 2021).

AlphaFold modelling generated 53,066 isoform-derived structures, of which 31,693 passed the mean-pLDDT threshold. These models were combined with 14,908 representative human reference structures, yielding an integrated structure set of 46,601 proteins.

#### Protein domains, intrinsic disorder and subcellular localization

Pfam domains were identified with HMMER hmmscan against Pfam-A v35.0 using curated trusted cut-offs (Eddy, 2011; Mistry et al., 2021). Redundant hits to the same Pfam family within a protein were resolved by retaining the highest-scoring compatible assignment. Intrinsically disordered regions were predicted from amino-acid sequences with IUPred3, and disorder content was summarized as total disordered length and fraction of the protein sequence (Erdős et al., 2021). Subcellular localization was predicted with DeepLoc 2.0 using the same protein-sequence set (Thumuluri et al., 2022). Domain, disorder and localization differences were evaluated between each non-reference proteoform and its corresponding reference proteoform.

### Structural motif analysis and benchmarking

#### Motif and zinc-finger annotation

Reviewed human UniProtKB/Swiss-Prot records, including canonical entries and annotated isoform sequence records, were downloaded on 8 August 2025. A total of 52,582 retrieved sequence records were processed; this number refers to downloaded sequence records rather than the number of canonical human genes. Residue-level annotations were extracted for motifs, zinc fingers, domains, binding sites, active sites and other curated functional features. Multiline annotations were normalized, and redundant or overlapping intervals of the same type within a protein were collapsed. UniProt and Ensembl identifiers were resolved against 3DisoGalaxy node identifiers to retain residue-level provenance.

Structural models used for motif analysis comprised quality-controlled AlphaFold Database references and in-house AlphaFold predictions. Models were normalized to single-chain, residue-indexed representations with validated sequence-to-structure correspondence. For AlphaFold Database proteins distributed across overlapping fragments, UniProt coordinates were converted to fragment-local coordinates using the fragment definitions supplied with the database models.

#### Folddisco search

Structure-based motif searches were conducted with Folddisco v2.0 on a SLURM-managed high-performance-computing cluster using 112 parallel workers (Kim et al., 2026). For each query protein, the structure fragment corresponding to the annotated motif interval was searched against the indexed 3DisoGalaxy structure set. Alignments were ranked using Folddisco geometric-fit and rarity-based scores. Search status and summary statistics were recorded for every query.

#### Benchmark definition and operating thresholds

Reviewed UniProtKB/Swiss-Prot functional and active-site annotations were used as the benchmark set. Zinc-finger intervals were obtained from the UniProt zinc-finger feature, and other functional motifs were obtained from residue-level feature annotations. The benchmark was restricted to proteins represented in the 3DisoGalaxy network. Candidate hits were initially screened with an RMSD cut-off of 3.0 Å and classified as follows: Tier A required RMSD ≤ 1.5 Å, coverage ≥ 0.80, gap ratio ≤ 0.05 and inverse document frequency score ≥ 0.40; Tier B required RMSD ≤ 2.5 Å, coverage ≥ 0.60, gap ratio ≤ 0.15 and inverse document frequency score ≥ 0.20.

Overlapping predictions were clustered within ±1 residue, and the highest-ranked representative was retained from each cluster. A retrieved region was considered a true positive when its mapped target interval overlapped a curated UniProt motif-bearing interval within a ±2-residue tolerance. The recovery benchmark was source-inclusive; therefore, the source protein from which a motif query was derived could also be present in the search database. Accordingly, this analysis was used as a pipeline-recovery benchmark rather than as an estimate of cross-protein motif-transfer performance. Non-zinc-finger analyses were stratified according to intrinsically disordered region (IDR) thresholds derived from IUPred3 (>0%, ≥50% or 100%); zinc-finger analyses were not filtered by this criterion. Prediction-level precision–recall ranking and average precision were calculated among clustered retrieved candidate regions. Uncertainty was evaluated with 200 target-level bootstrap replicates, each containing 70% of eligible target proteins. Paired tests were used when comparing IDR thresholds within the same bootstrap samples.

False-discovery behaviour was evaluated with two decoy sets: hard-negative intervals that did not overlap benchmark sites or known zinc-finger signatures, and length- and quality-matched random intervals sampled from the same target proteins while excluding benchmark and hard-negative regions. Estimated false-discovery rates were calculated from decoy hits relative to the corresponding true-positive detections at each operating point. For zero-decoy-hit settings, a conservative 95% upper bound was estimated using the rule of three. Tier A matches satisfying the IDR ≥ 50% criterion were used for downstream general-motif analyses; Tier A matches without an IDR filter were used for zinc-finger analyses.

#### Motif gain, loss and cancer-driver comparisons

Motif remodelling was quantified at the gene or isoform-family level. For each gene, the reference motif set, A, was defined as the union of motif keys across its reference isoforms, and the non-reference set, B, as the union across its non-reference isoforms. Genes were included when both sets were represented and their union was non-empty. Motif gain and loss were defined as |B \ A| and |A \ B|, respectively. Motif-set similarity was measured with the Jaccard index, J = |A ∩ B|/|A ∪ B|, and dissimilarity was expressed as 1 − J. Gain and loss counts were compared across genes with a two-sided Wilcoxon signed-rank test after excluding ties, with a two-sided exact binomial sign test as a complementary analysis.

Cancer-driver status was assigned using the IntOGen catalogue (Martínez-Jiménez et al., 2020). A motif-shift score was calculated for each gene as (gain + loss) + (1 − J), corresponding to a Jaccard-distance weight of 1.0. Score distributions in cancer-driver and non-driver genes were compared with a two-sided Mann–Whitney U test. Violin plots were overlaid with boxplots showing medians and interquartile ranges.

### Structural-similarity network

All-versus-all structural searches were performed with Foldseek easy-search using global TM-align-type alignment (--alignment-type 1; --tmalign-fast 1) (van Kempen et al., 2024). Proteoforms were represented as network nodes, and pairs with TM-score ≥ 0.9 were retained as edges. Connected components of this thresholded graph were treated as structural clusters. For clusters containing at least two nodes with Pfam annotations, domain consistency was summarized as the mean pairwise Jaccard index between Pfam-accession sets. Structural summaries included mean TM-score and alignment lDDT.

Network visualization was performed in Gephi v0.10.1 using ForceAtlas2 with 31 threads, tolerance 1.0, approximation 1.2, scaling 10.0, gravity 0.1, stronger gravity enabled, hub dissuasion enabled, LinLog mode disabled, overlap prevention enabled and edge-weight influence set to 1.0 (Jacomy et al., 2014). Protein structures and residue-level motif mappings were rendered in PyMOL v3.1.6.1.

### 3DisoGalaxy web portal

The 3DisoGalaxy portal was implemented as an interactive interface for querying the proteoform structural-similarity network. Network rendering and interaction were implemented with Cosmograph v1.3.0. Nodes can be searched by gene symbol or protein identifier, and sequence- or structure-based searches use Foldseek to retrieve structurally related proteoforms. Three-dimensional structures are displayed with Mol* v3.35.0 (Sehnal et al., 2021). Node metadata include transcript and protein identifiers, tissue or subtype provenance, expression attributes and local structural-neighbourhood information.

### Experimental validation

#### Cell culture

MCF-10A human mammary epithelial cells and ZR-75-1, BT-474, HCC38, HCC1806 and MDA-MB-231 breast cancer cells were obtained from ATCC. MCF-10A cells were maintained in specialized medium (Procell, CM-0525). ZR-75-1, BT-474, HCC38 and HCC1806 cells were maintained in RPMI-1640 medium (Gibco, 11875-093) supplemented with 10% fetal bovine serum (Gibco, 10099-141) and 1% penicillin–streptomycin (Gibco, 15140-122) at 37 °C in a humidified 5% CO2 atmosphere. MDA-MB-231 cells were maintained in Leibovitz’s L-15 medium (Gibco, 41300-039) supplemented with 10% fetal bovine serum and 1% penicillin–streptomycin at 37 °C without additional CO2.

#### RNA extraction, reverse transcription and RT-PCR

Total RNA was extracted with the SteadyPure Universal RNA Extraction Kit (Accurate Biotechnology, AG21022). RNA concentration and purity were evaluated with a NanoDrop 2000 spectrophotometer, and samples with A260/A280 ratios between 1.8 and 2.0 were used for reverse transcription. cDNA was synthesized from 1 μg total RNA with HiScript III RT SuperMix for qPCR (Vazyme, R323-01) using 25 °C for 5 min, 50 °C for 15 min and 85 °C for 5 s, followed by a 4 °C hold.

Primer pairs were designed to distinguish products predicted from canonical and candidate non-canonical transcript models. PCR was performed in 20 μl reactions containing 10 μl 2× Taq PCR Master Mix (Vazyme, P112-02), 0.4 μM of each primer and 1 μl cDNA. Cycling conditions were 95 °C for 5 min; 35 cycles of 95 °C for 30 s, 55 °C for 30 s and 72 °C for 30 s; and 72 °C for 5 min. Products were separated on 3% agarose gels, stained with GelRed (Biotium, 41003) and visualized with a gel-imaging system. β-actin was used as an internal control.

The recorded primer sequences were as follows: HLA-C canonical forward 5′- ACCTGCGCGGCTACTACAAC-3′, candidate non-canonical forward 5′- CGGGTATGACCAGTCCGCCT-3′ and shared reverse 5′-CTGCAGCGTCTCCTTCCCG-3′; SLC4A2 canonical forward 5′-GACCAGATCAAGGCCGAGG-3′, candidate non-canonical forward 5′- GCCACGGTGGTCCTTGTGG-3′ and shared reverse 5′-GCCTCGTGGAATTGCTTGTC-3′; and SLC39A1 recorded shared forward 5′-TTCCTGGTCCTGGTGATGGAGC-3′ and shared reverse 5′- GGCCCACCATTCACTGTTC-3′.

#### Sanger sequencing of RT-PCR products

The lower, candidate non-canonical-sized HLA-C, SLC4A2 and SLC39A1 RT-PCR products were subjected to Sanger sequencing. Chromatogram traces were quality-checked from the original AB1 files and compared with the corresponding predicted amplicon sequences. The SLC4A2 read crossed the predicted novel exon–exon junction. The HLA-C and SLC39A1 reads confirmed the identities of their respective amplicons but did not independently resolve transcript-specific splice junctions.

#### Protein extraction and immunoblotting

Total protein was extracted in RIPA buffer (Beyotime, P0013B) supplemented with protease-inhibitor cocktail (Roche, 04693159001) and phosphatase inhibitors (Beyotime, P1081). Cells were lysed on ice for 30 min with mixing every 10 min and centrifuged at 12,000 × g for 15 min at 4 °C. Protein concentrations in clarified lysates were measured with the BCA Protein Assay Kit (Beyotime, P0012).

The canonical AKT1 sequence comprises 480 amino acids and has a calculated molecular mass of approximately 55.7 kDa. The TransDecoder-selected AKT1-ΔPH candidate used for structure prediction comprises 334 amino acids corresponding to residues 147–480 of canonical AKT1 and has a calculated molecular mass of approximately 38.6 kDa. Twenty micrograms of total protein per cell line were separated by 4–16% SDS–PAGE and transferred to PVDF membranes. Membranes were probed with an AKT1 antibody (Proteintech, 80457-1-RR); β-actin (Proteintech, 66009-1-Ig) was used as the loading control. Immunoreactive bands were detected by enhanced chemiluminescence and quantified with ImageJ (Schneider et al., 2012). A lower-molecular-weight AKT1-reactive band observed in addition to the canonical-size band was selected for LC–MS/MS analysis.

#### LC–MS/MS analysis of AKT1-reactive gel regions

For proteomic characterization, 30–50 μg total protein was separated by SDS–PAGE, and gel regions corresponding to the canonical-size and lower-molecular-weight AKT1-reactive species were excised. Samples were analysed by Shanghai Fuming Biotechnology. Parallel in-gel digestion was performed with trypsin, Lys-C, Lys-N, Asp-N, chymotrypsin and Arg-C. LC–MS/MS data were processed with Proteome Discoverer v2.5 and searched against the UniProt human protein database supplemented with the predicted 334-amino-acid AKT1-ΔPH sequence.

AKT1-derived peptides detected in canonical-size and lower-molecular-weight gel regions were mapped to the canonical AKT1 sequence and to the sequence retained in the AKT1-ΔPH candidate. Peptide-spectrum matches and sequence coverage were evaluated for each gel region. Because peptides from the lower-molecular-weight region mapped to sequence shared by canonical AKT1 and the predicted candidate, these data support the AKT1 identity of the lower band but do not independently establish its complete sequence, precise N terminus or identity as the intact predicted AKT1-ΔPH proteoform.

## Data availability

All data required to evaluate the conclusions of this study are provided within the paper and its Supplementary Information. Curated datasets and additional materials are publicly available via Zenodo (DOI: 10.5281/zenodo.14865747). Model predictions generated in this study are accessible through an interactive online platform at http://3disogalaxy.com/, with accompanying documentation available at https://github.com/FeliciaTJiang/3DisoGalaxy.

## Code availability

Source code, analysis R and python scripts for bioinformatic analysis are available at https://github.com/FeliciaTJiang/3DisoGalaxy.

## References

Aebersold, R. & Mann, M. Mass-spectrometric exploration of proteome structure and function. Nature 537, 347–355 (2016).

Aebersold, R. et al. How many human proteoforms are there? Nat. Chem. Biol. 14, 206–214 (2018).

Akdel, M. et al. A structural biology community assessment of AlphaFold2 applications. Nat. Struct. Mol. Biol. 29, 1056–1067 (2022).

Alamancos, G. P., Pagès, A., Trincado, J. L., Bellora, N. & Eyras, E. Leveraging transcript quantification for fast computation of alternative splicing profiles. RNA 21, 1521–1531 (2015).

Buljan, M. et al. Tissue-specific splicing of disordered segments that embed binding motifs rewires protein interaction networks. Mol. Cell 46, 871–883 (2012).

Camacho, C., et al. BLAST+: architecture and applications. BMC Bioinformatics 10, 421 (2009).

Carpten, J. D. et al. A transforming mutation in the pleckstrin homology domain of AKT1 in cancer. Nature 448, 439–444 (2007).

Danecek, P. et al. Twelve years of SAMtools and BCFtools. GigaScience 10, giab008 (2021).

Dobin, A. et al. STAR: ultrafast universal RNA-seq aligner. Bioinformatics 29, 15–21 (2013).

Dvinge, H., Kim, E., Abdel-Wahab, O. & Bradley, R. K. RNA splicing factors as oncoproteins and tumour suppressors. Nat. Rev. Cancer 16, 413–430 (2016).

Eddy, S. R. Accelerated profile HMM searches. PLoS Comput. Biol. 7, e1002195 (2011).

Ellis, J. D. et al. Tissue-specific alternative splicing remodels protein–protein interaction networks. Mol. Cell 46, 884–892 (2012).

Erdős, G., Pajkos, M. & Dosztányi, Z. IUPred3: prediction of protein disorder enhanced with unambiguous experimental annotation and visualization of evolutionary conservation. Nucleic Acids Res. 49, W297–W303 (2021).

Ewels, P., Magnusson, M., Lundin, S. & Käller, M. MultiQC: summarize analysis results for multiple tools and samples in a single report. Bioinformatics 32, 3047–3048 (2016).

Ezkurdia, I. et al. Most highly expressed protein-coding genes have a single dominant isoform. J. Proteome Res. 14, 1880–1887 (2015).

Frankish, A. et al. GENCODE: reference annotation for the human and mouse genomes in 2023. Nucleic Acids Res. 51, D942–D949 (2023).

Glinos, D. A. et al. Transcriptome variation in human tissues revealed by long-read sequencing. Nature 608, 353–359 (2022).

Holehouse, A. S. & Kragelund, B. B. The molecular basis for cellular function of intrinsically disordered protein regions. Nat. Rev. Mol. Cell Biol. 25, 187–211 (2024).

Ingolia, N. T., Ghaemmaghami, S., Newman, J. R. S. & Weissman, J. S. Genome-wide analysis in vivo of translation with nucleotide resolution using ribosome profiling. Science 324, 218–223 (2009).

Ingolia, N. T., Hussmann, J. A. & Weissman, J. S. Ribosome profiling: global views of translation. Cold Spring Harb. Perspect. Biol. 11, a032698 (2019).

Jacomy, M., Venturini, T., Heymann, S. & Bastian, M. ForceAtlas2, a continuous graph layout algorithm for handy network visualization designed for the Gephi software. PLoS ONE 9, e98679 (2014).

Jiang, F. T. et al. Structure-aware protein function prediction at isoform resolution. bioRxiv 10.64898/2026.04.24.720502 (2026).

Jumper, J. et al. Highly accurate protein structure prediction with AlphaFold. Nature 596, 583–589 (2021).

Kahles, A. et al. Comprehensive analysis of alternative splicing across tumors from 8,705 patients. Cancer Cell 34, 211–224.e6 (2018).

Kim, H. et al. Structural motif search across the protein universe with Folddisco. Nat. Biotechnol. 10.1038/s41587-026-03162-9 (2026).

Langmead, B., Trapnell, C., Pop, M. & Salzberg, S. L. Ultrafast and memory-efficient alignment of short DNA sequences to the human genome. Genome Biol. 10, R25 (2009).

Lau, E. et al. Splice-junction-based mapping of alternative isoforms in the human proteome. Cell Rep. 29, 3751–3765.e5 (2019).

Li, H. Minimap2: pairwise alignment for nucleotide sequences. Bioinformatics 34, 3094–3100 (2018).

Liu, Y. et al. Impact of alternative splicing on the human proteome. Cell Rep. 20, 1229–1241 (2017).

Manning, B. D. & Toker, A. AKT/PKB signaling: navigating the network. Cell 169, 381–405 (2017).

Martin, M. Cutadapt removes adapter sequences from high-throughput sequencing reads. EMBnet.journal 17, 10–12 (2011).

Martínez-Jiménez, F. et al. A compendium of mutational cancer driver genes. Nat. Rev. Cancer 20, 555–572 (2020).

Mirdita, M. et al. ColabFold: making protein folding accessible to all. Nat. Methods 19, 679–682 (2022).

Mistry, J. et al. Pfam: the protein families database in 2021. Nucleic Acids Res. 49, D412–D419 (2021).

Morales, J. et al. A joint NCBI and EMBL-EBI transcript set for clinical genomics and research. Nature 604, 310–315 (2022).

Nguyen Ba, A. N., et al. Proteome-wide discovery of evolutionary conserved sequences in disordered regions. Sci. Signal. 5, rs1 (2012).

Pan, Q., Shai, O., Lee, L. J., Frey, B. J. & Blencowe, B. J. Deep surveying of alternative splicing complexity in the human transcriptome by high-throughput sequencing. Nat. Genet. 40, 1413–1415 (2008).

Pardo-Palacios, F. J. et al. SQANTI3: curation of long-read transcriptomes for accurate identification of known and novel isoforms. Nat. Methods 21, 793–797 (2024).

Reixachs-Solé, M., Ruiz-Orera, J., Albà, M. M. & Eyras, E. Ribosome profiling at isoform level reveals evolutionary conserved impacts of differential splicing on the proteome. Nat. Commun. 11, 1768 (2020).

Schneider, C. A., Rasband, W. S. & Eliceiri, K. W. NIH Image to ImageJ: 25 years of image analysis. Nat. Methods 9, 671–675 (2012).

Sehnal, D. et al. Mol* Viewer: modern web app for 3D visualization and analysis of large biomolecular structures. Nucleic Acids Res. 49, W431–W437 (2021).

Sharon, D., Tilgner, H., Grubert, F. & Snyder, M. A single-molecule long-read survey of the human transcriptome. Nat. Biotechnol. 31, 1009–1014 (2013).

Smith, L. M., Kelleher, N. L. & The Consortium for Top Down Proteomics. Proteoform: a single term describing protein complexity. Nat. Methods 10, 186–187 (2013).

The UniProt Consortium. UniProt: the Universal Protein Knowledgebase in 2023. Nucleic Acids Res. 51, D523–D531 (2023).

Thul, P. J. et al. A subcellular map of the human proteome. Science 356, eaal3321 (2017).

Thumuluri, V. et al. DeepLoc 2.0: multi-label subcellular localization prediction using protein language models. Nucleic Acids Res. 50, W228–W234 (2022).

Tran, J. C. et al. Mapping intact protein isoforms in discovery mode using top-down proteomics. Nature 480, 254–258 (2011).

Trincado, J. L. et al. SUPPA2: fast, accurate, and uncertainty-aware differential splicing analysis across multiple conditions. Genome Biol. 19, 40 (2018).

Tunyasuvunakool, K. et al. Highly accurate protein structure prediction for the human proteome. Nature 596, 590–596 (2021).

van Kempen, M. et al. Fast and accurate protein structure search with Foldseek. Nat. Biotechnol. 42, 243–246 (2024).

Van Roey, K. et al. Short linear motifs: ubiquitous and functionally diverse protein interaction modules directing cell regulation. Chem. Rev. 114, 6733–6778 (2014).

Varadi, M. et al. AlphaFold Protein Structure Database: massively expanding the structural coverage of protein-sequence space with high-accuracy models. Nucleic Acids Res. 50, D439–D444 (2022).

Wang, E. T. et al. Alternative isoform regulation in human tissue transcriptomes. Nature 456, 470–476 (2008).

Whitley, M. J. et al. Comparative analysis of KRAS4a and KRAS4b splice variants reveals distinctive structural and functional properties. Sci. Adv. 10, eadj4137 (2024).

Wright, P. E. & Dyson, H. J. Intrinsically disordered proteins in cellular signalling and regulation. Nat. Rev. Mol. Cell Biol. 16, 18–29 (2015).

Wyman, D. et al. A technology-agnostic long-read analysis pipeline for transcriptome discovery and quantification. bioRxiv 10.1101/672931 (2020).

Xiao, Z. et al. De novo annotation and characterization of the translatome with ribosome profiling data. Nucleic Acids Res. 46, e61 (2018).

Yang, X. et al. Widespread expansion of protein interaction capabilities by alternative splicing. Cell 164, 805–817 (2016).

