## Supplementary material for "Alternative splicing expands and remodels the breast cancer proteoform landscape": Supplemnentary figures

### 3DisoGalaxy portal integrates the transcriptome, translatome and foldome

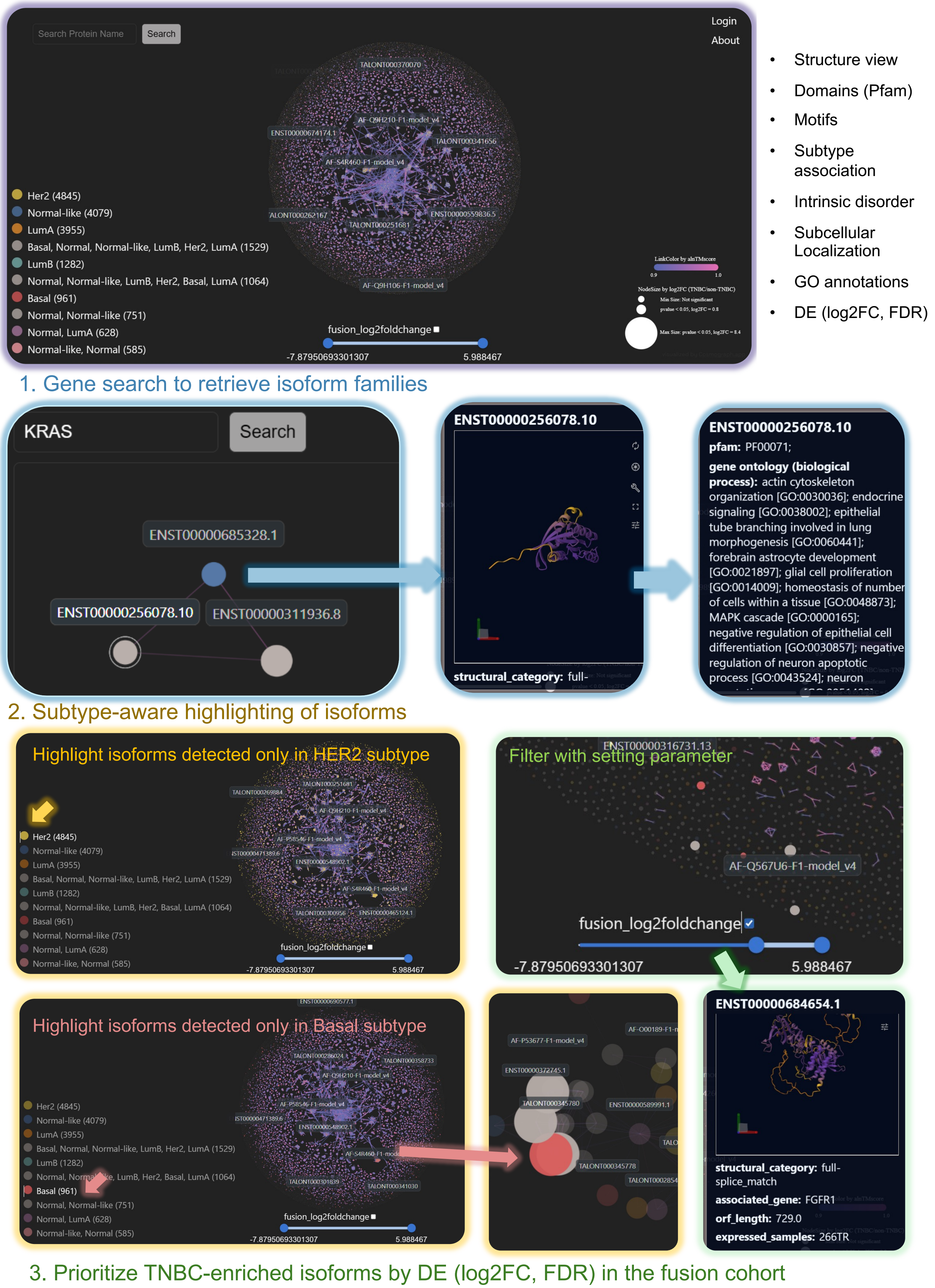


**Supplementary Fig. 1.** 3DisoGalaxy portal for isoform-resolved exploration of the breast-cancer transcriptome, translatome and foldome in a structure-similarity network (3disogalaxy.com).

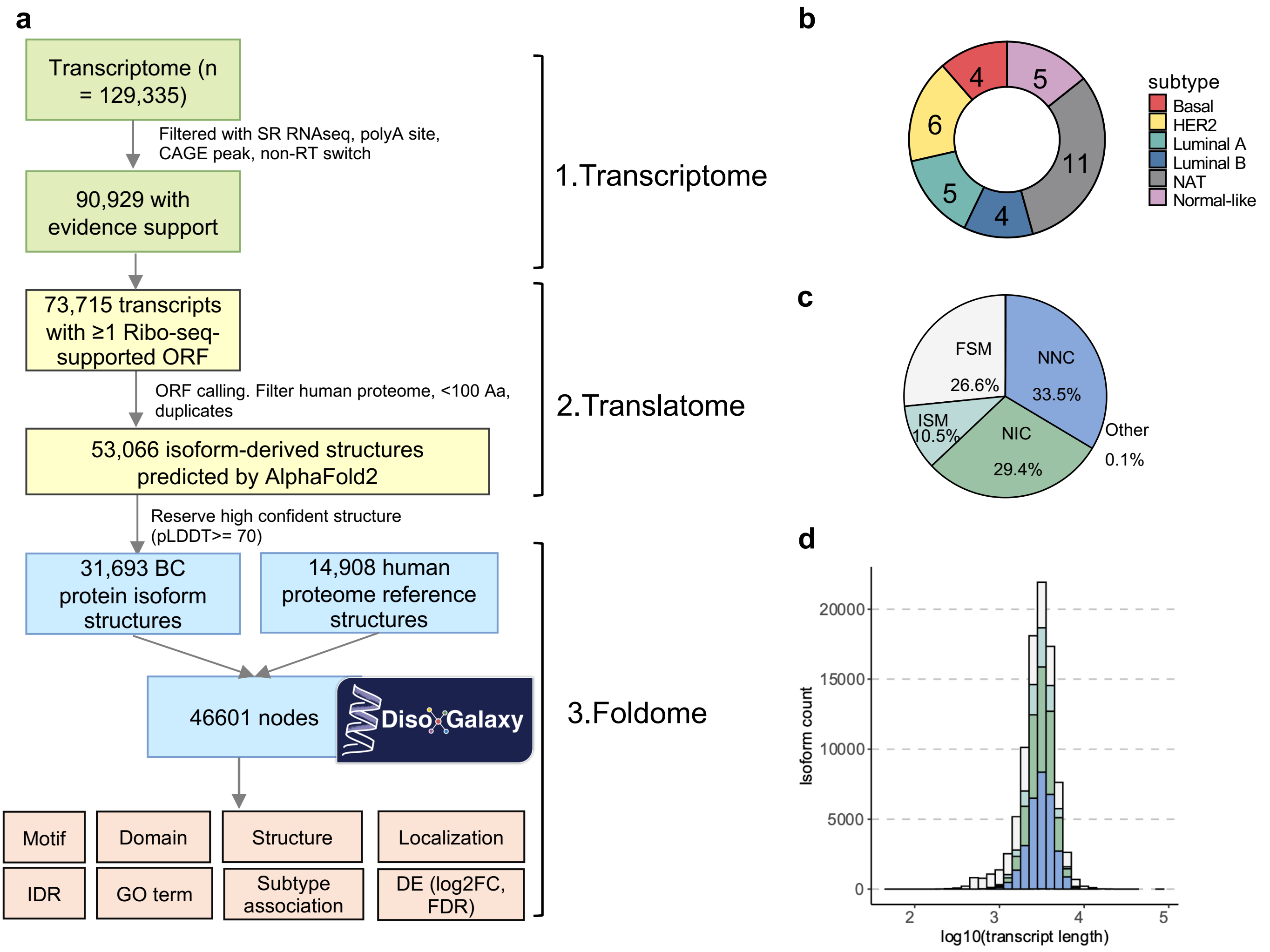


**Supplementary Fig. 2.** Cohort overview, evidence curation, and basic properties of the breast-cancer transcript-variant set. **(A)** Workflow from transcript discovery and evidence curation to ORF definition, translation support, and foldome construction. **(B)** Subtype composition of the profiled cohort. **(C)** Transcript-variant distribution across reference-guided structural categories. **(D)** Transcript-length distributions (log10 scale) for the curated transcript-variant set.

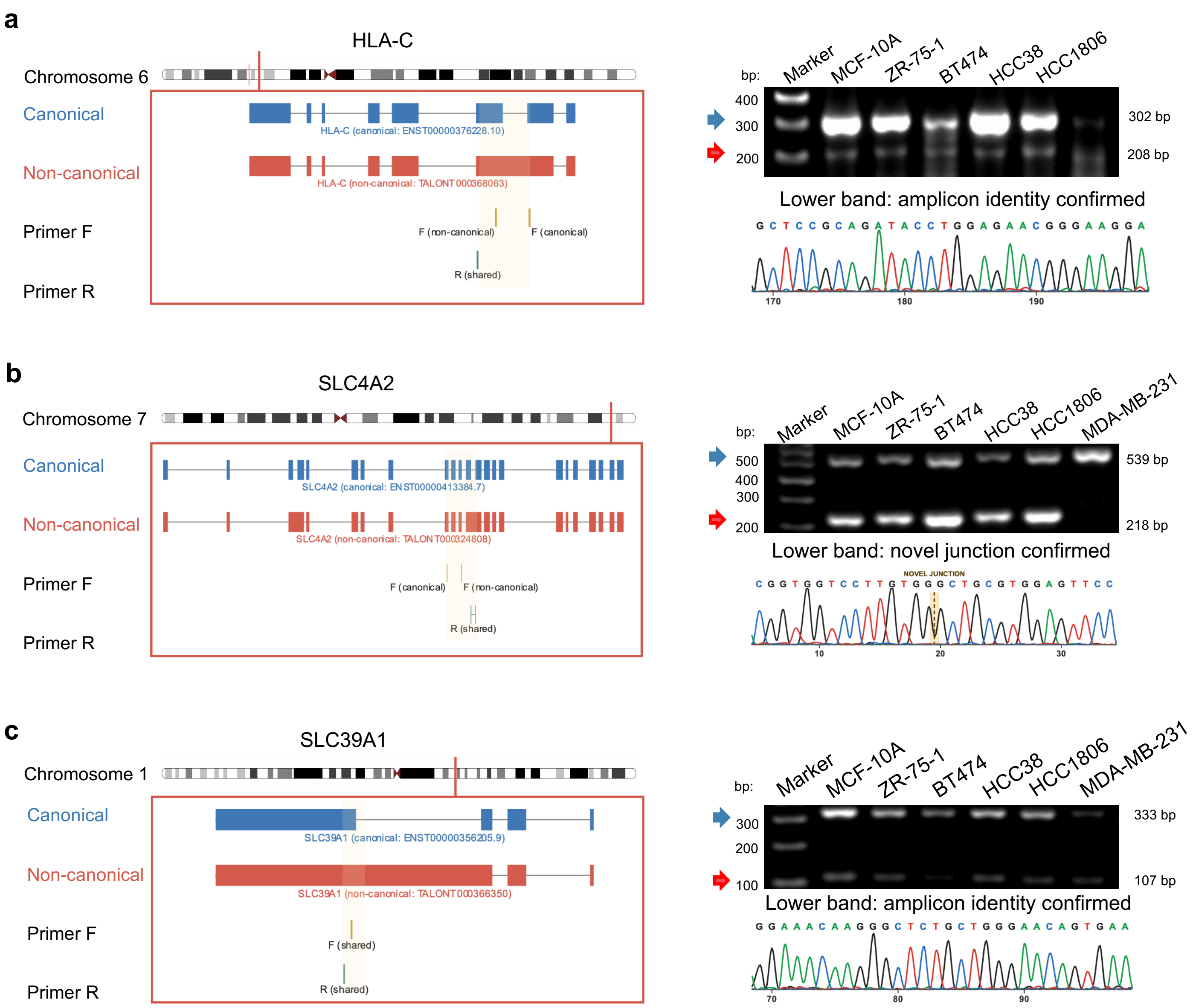


**Supplementary Fig. 3 | Experimental validation of candidate non-canonical transcript models by RT–PCR and Sanger sequencing.** Transcript structures and primer positions are shown for HLA-C, SLC4A2 and SLC39A1. RT–PCR products of the expected sizes supported the expression of the canonical and candidate non-canonical transcript models identified by 3DisoGalaxy. Blue and red arrows indicate the predicted canonical and non-canonical products, respectively. The non-canonical-sized products from HLA-C, SLC4A2 and SLC39A1 were subjected to Sanger sequencing. The SLC4A2 sequence read directly spanned the predicted novel exon–exon junction, whereas the HLA-C and SLC39A1 reads confirmed the identities of the corresponding amplicons but did not independently resolve transcript-specific splice junctions. Lanes correspond to the indicated breast cancer and normal mammary cell lines.

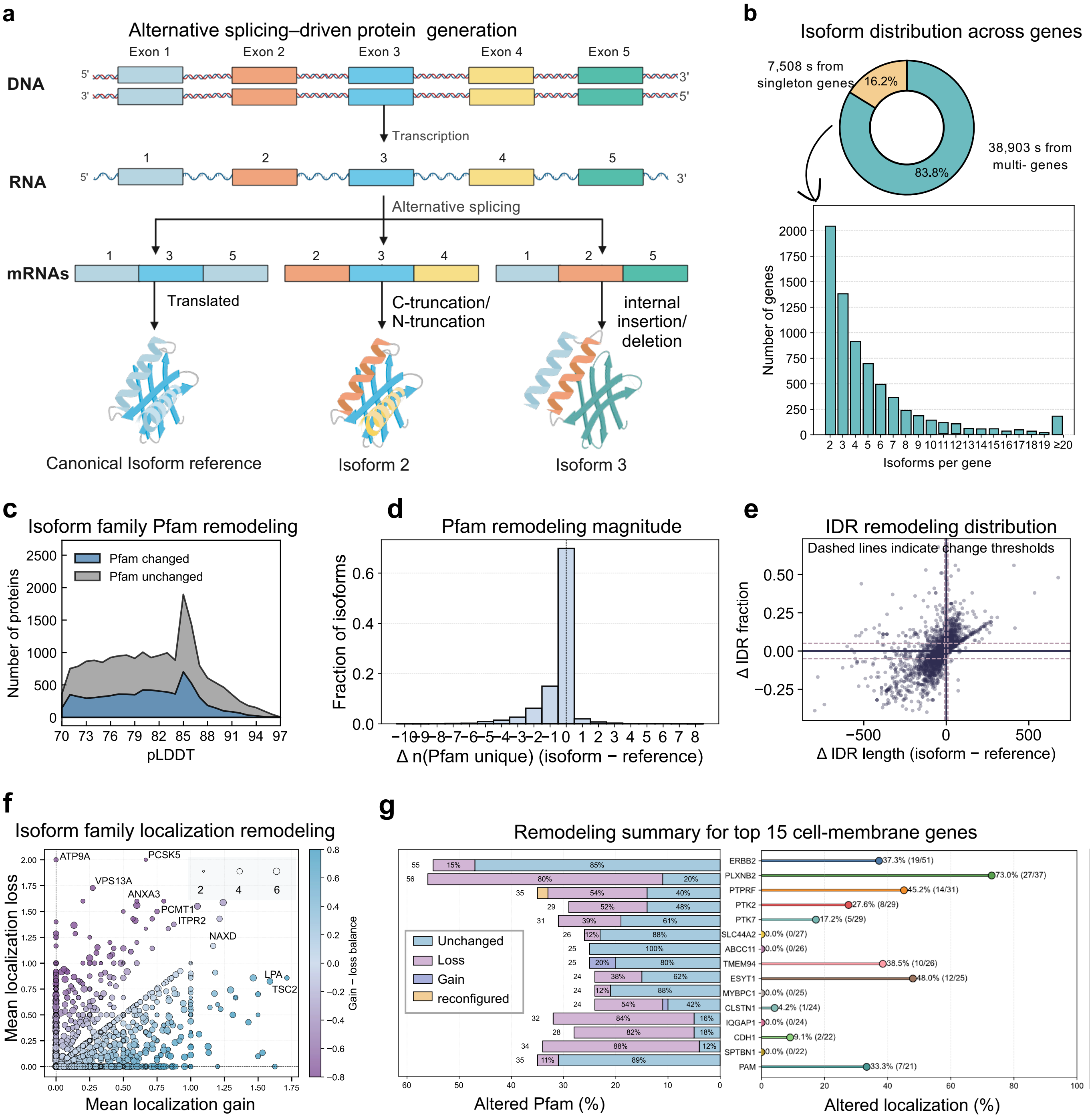


**Supplementary Fig. 4.** Isoform-family remodeling of Pfam domains, intrinsic disordered regions (IDRs), and predicted localization. **(a)** Alternative splicing generates protein isoforms with C- or N-terminal truncations and internal insertions or deletions relative to the canonical reference isoform. **(b)** Distribution of isoform numbers across genes, showing singleton and multi-isoform genes. **(c)** pLDDT distributions for isoforms with or without Pfam remodeling within isoform families. **(d)** Change in unique Pfam domain count for each isoform relative to its reference isoform. **(e)** Isoform-reference differences in IDR length and IDR fraction, with dashed lines indicating remodeling thresholds. **(f)** Isoform-family-level predicted localization remodeling, summarized by mean localization gain and loss. **(g)** Top 15 isoform-rich cell-membrane genes ranked by Pfam remodeling composition and altered predicted localization fraction.

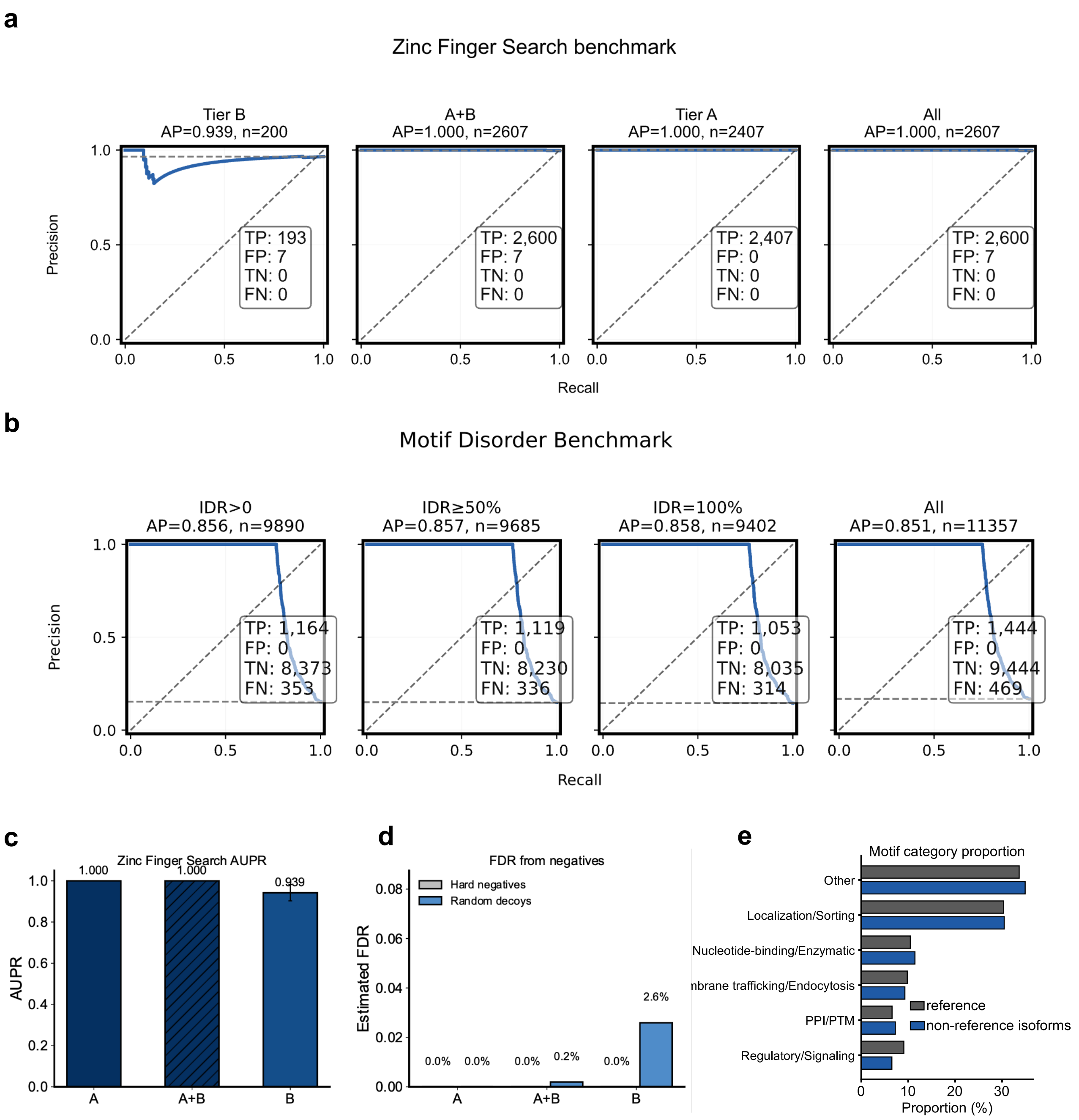


**Supplementary Fig. 5. Benchmarking and controls for structure-based motif and zinc-finger detection. a,** Prediction-level precision–recall ranking of retrieved zinc-finger structural matches under tiered structural-match stringency settings (Tier B, combined Tier A and B, Tier A and pooled), with average precision (AP) and sample size shown for each set. **b,** Prediction-level precision–recall ranking of retrieved non-zinc-finger structural matches stratified by intrinsically disordered region (IDR) thresholds (IDR > 0, IDR ≥ 50% and IDR = 100%) and for the pooled set, with average precision (AP) and sample size shown for each set. **c,** Summary of prediction-level average precision for zinc-finger structural matches across Tier A, combined Tier A and B, and Tier B. **d,** Decoy-based false-discovery assessment for zinc-finger detection using hard-negative intervals and random decoys, reported as estimated false-discovery rates for each operating point. **e,** Distribution of putative motif-category assignments in reference and non-reference isoforms.

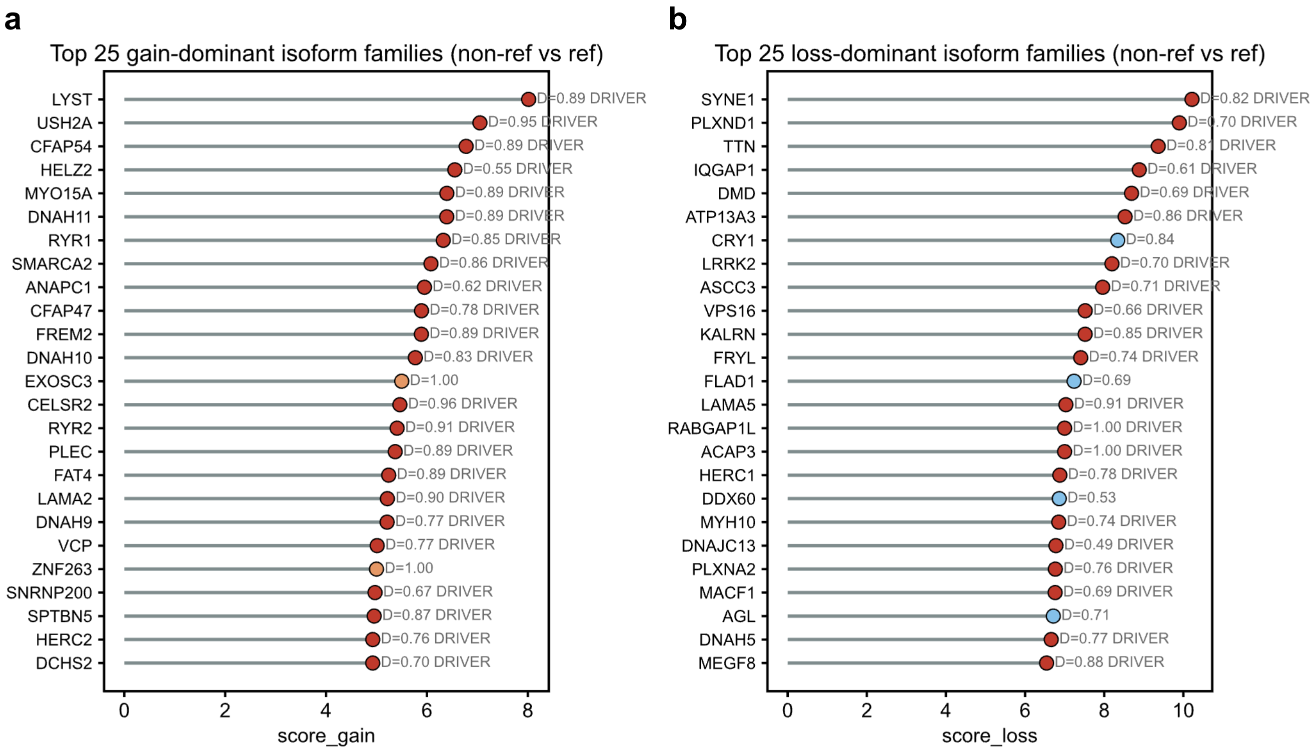


**Supplementary Fig. 6**. Top gene-level isoform families with gain- or loss-dominant pooled motif repertoires. **a.** Top 25 gene-level isoform families ranked by motif gain score for the pooled non-reference versus reference repertoire. **b.** Top 25 gene-level isoform families ranked by motif loss score for the pooled non-reference versus reference repertoire.

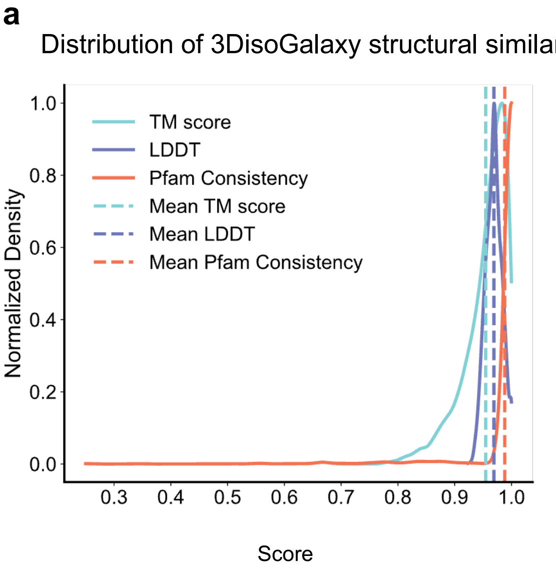


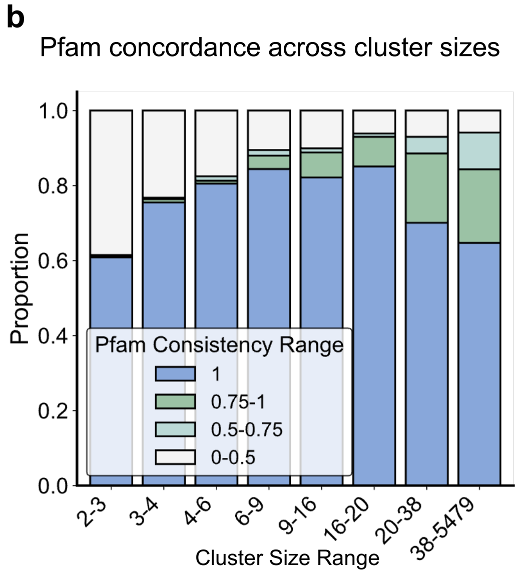


**Supplementary Fig. 7. Global structural similarity and Pfam concordance in the 3DisoGalaxy network. a,** Normalized density distributions of TM score, mean pLDDT and Pfam concordance across structure-similarity edges; dashed lines indicate the corresponding mean values. **b,** Proportional composition of four Pfam-concordance intervals across non-singleton cluster-size ranges.
